# Discovery of a brain-penetrant CHI3L1 inhibitor as an oral modulator of neuroinflammation in Alzheimer’s disease

**DOI:** 10.64898/2026.10.15.759511

**Authors:** Saurabh Upadhyay, Longfei Zhang, Baljit Kaur, Laura Calvo-Barreiro, Marianna Teplova, Moustafa T. Gabr

## Abstract

Chitinase-3-like protein 1 (CHI3L1) is elevated in Alzheimer’s disease (AD), but its role in glial dysfunction and therapeutic tractability remain unclear. Here, screening a 4.2-billion-compound DNA-encoded library identified DELC2, a brain-penetrant CHI3L1 modulator. Biophysical and crystallographic studies showed that DELC2 binds the CHI3L1 glycan-recognition groove within an aromatic-hydrophobic pocket centered on Trp99 and Trp352. DELC2 inhibited CHI3L1-RAGE association, suppressed CHI3L1-induced inflammatory signaling, reduced Gal-3-positive microglia, and restored amyloid-β uptake in glial cultures. In human cerebrospinal fluid, CHI3L1 was elevated in AD and correlated with astrocytic NF-κB activation. DELC2 phenocopied CHI3L1 immunodepletion, whereas CHI3L1 add-back restored inflammatory and amyloid-β-handling defects. Oral DELC2 achieved sustained unbound brain exposure in 5xFAD mice, improved cognitive performance, reduced multiple amyloid-β species, and attenuated astrocytic and microglial inflammatory programs. These findings establish the CHI3L1 glycan-recognition groove as a druggable regulator of RAGE-associated glial dysfunction and identify DELC2 as a chemical scaffold for CHI3L1-directed therapy in AD.

## Introduction

Alzheimer’s disease (AD) is a progressive neurodegenerative disorder characterized by cognitive decline, synaptic dysfunction, and irreversible neuronal loss, affecting millions of individuals worldwide (1). Although extracellular amyloid-β (Aβ) plaques and intracellular neurofibrillary tangles composed of hyperphosphorylated tau remain defining pathological hallmarks of the disease, increasing evidence indicates that chronic neuroinflammation is a central driver of disease progression rather than a secondary consequence of protein aggregation alone (2). Persistent activation of astrocytes and microglia establishes a self-reinforcing inflammatory environment that exacerbates synaptic dysfunction, impairs neuronal repair mechanisms, and accelerates neurodegeneration (3). Reactive astrocytes lose critical homeostatic functions, including metabolic support, regulation of synaptic activity, maintenance of extracellular ion balance, and clearance of toxic protein aggregates, thereby contributing directly to disease pathology (4). In parallel, inflammatory crosstalk between astrocytes and microglia amplifies cytokine production, oxidative stress, and defective phagocytic responses, ultimately impairing Aβ clearance and neuronal homeostasis (5, 6).

Among the astrocyte-derived inflammatory mediators implicated in AD progression, chitinase-3-like protein 1 (CHI3L1), also known as YKL-40, has emerged as both a robust biomarker and an active pathogenic regulator of neuroinflammatory signaling (7, 8). CHI3L1 is a secreted glycoprotein predominantly produced by reactive astrocytes, and elevated levels in cerebrospinal fluid (CSF) and plasma strongly correlate with tau pathology, cortical atrophy, cognitive decline, and progression from presymptomatic to symptomatic stages of AD (9, 10). Histopathological studies further demonstrate that CHI3L1 expression is enriched in astrocytes localized near amyloid plaques and tau-positive regions, linking its expression to sites of active neurodegeneration (11, 12). Importantly, recent functional studies indicate that CHI3L1 is not merely a passive biomarker of glial activation but instead acts as a signaling mediator that directly modulates astrocytic and neuronal function (13). Astrocyte-specific deletion of CHI3L1 in AD mouse models reduces amyloid burden, suppresses microgliosis, restores synaptic integrity, and improves cognitive performance, establishing CHI3L1 as a causal effector linking reactive astrocytes to neurodegenerative progression (14).

Mechanistically, CHI3L1 participates in multiple receptor-mediated signaling pathways that converge on neuroinflammatory and neurodegenerative dysfunction (15). Previous studies demonstrated that astrocyte-secreted CHI3L1 engages the receptor for advanced glycation end products (RAGE), activating NF-κB-dependent inflammatory signaling and promoting reactive astrocyte phenotypes associated with impaired glial homeostasis (7). In neurons, CHI3L1–RAGE signaling induces GSK3β-mediated destabilization of β-catenin, resulting in dendritic simplification, synaptic dysfunction, and neuronal injury (16). In parallel, CHI3L1 has been shown to activate CRTH2-dependent signaling in hippocampal neural stem cells, triggering the IKKβ–S6K1–S6 axis and suppressing neurogenesis (17). Additional evidence indicates that CHI3L1 negatively regulates glial phagocytic and lysosomal activity, thereby impairing Aβ uptake and clearance while reinforcing chronic inflammatory signaling (18). Collectively, these findings position CHI3L1 as a central neuroimmune signaling node linking astrocyte activation, impaired amyloid handling, inflammatory amplification, and neuronal dysfunction in AD (19).

Despite compelling biological and translational evidence implicating CHI3L1 in neurodegeneration, therapeutic targeting of this protein remains underdeveloped. Antibody-and oligonucleotide-based approaches are constrained by limited blood–brain barrier penetration, poor CNS exposure, immunogenicity concerns, and manufacturing complexity, limiting their suitability for chronic neurological diseases (20–22). By contrast, small molecules offer several advantages for targeting neuroinflammatory pathways, including oral bioavailability, tunable pharmacokinetic properties, scalable synthesis, and improved potential for CNS penetration (23, 24). However, CHI3L1 has remained a particularly challenging target for small-molecule discovery. Unlike catalytically active chitinases, CHI3L1 lacks enzymatic activity due to substitutions within the catalytic motif, functioning instead as a pseudoenzymatic signaling glycoprotein (25). The CHI3L1 glycan-recognition groove is relatively shallow and conformationally dynamic, lacking the well-defined catalytic pockets typically exploited in structure-based drug discovery (26). Consequently, only a limited number of CHI3L1-binding small molecules have been reported, and the structural basis governing ligand recognition and pharmacological modulation of CHI3L1 remains poorly understood (27, 28).

DNA-encoded library (DEL) technology has emerged as a powerful platform for identifying ligands against challenging and unconventional protein targets, particularly extracellular proteins that lack robust activity-based screening assays (29). Because DEL screening relies on affinity-based enrichment rather than catalytic readouts, it is especially well suited for pseudoenzymatic proteins such as CHI3L1 (13). In addition, DEL screening enables interrogation of billions of structurally diverse compounds using minimal quantities of recombinant protein, thereby facilitating discovery of ligands against difficult extracellular recognition interfaces (30). We therefore hypothesized that DEL screening could identify structurally distinct ligands capable of engaging the dynamic glycan-recognition architecture of CHI3L1.

Here, we report the discovery and structural characterization of DELC2, a CHI3L1-targeted small molecule identified through DEL screening and validated using orthogonal biophysical, structural, and functional approaches. Direct interaction between DELC2 and recombinant human CHI3L1 was confirmed using microscale thermophoresis (MST) and surface plasmon resonance (SPR), demonstrating reproducible micromolar target engagement across orthogonal biophysical platforms. X-ray crystallographic analysis revealed that DELC2 occupies the canonical glycan-recognition groove of CHI3L1 through an aromatic-enclosure binding mode centered on the dynamic Trp99/Trp352 recognition network. Rather than functioning as a classical carbohydrate mimic, DELC2 stabilizes a compact hydrophobic groove architecture through extensive π-stacking and hydrophobic packing interactions, supporting a potential mechanism involving conformational restriction of the CHI3L1 recognition cleft. Finally, functional evaluation in AD-relevant cellular models demonstrated that DELC2 modulates neuroinflammatory signaling and Aβ-associated pathological responses. Together, these findings establish DELC2 as a structurally distinct CHI3L1-targeted scaffold and provide a structural and mechanistic framework for pharmacological modulation of astrocyte-driven neuroinflammatory signaling in Alzheimer’s disease.

## Methods

### DNA-Encoded Library (DEL) Screening

The DEL screening experiment was performed as previously described (13). Briefly, affinity selection was conducted using a DNA-encoded chemical library of 4.2 billion compounds (WuXi AppTec) against immobilized human CHI3L1 protein (Cat. # CH1-H5228, ACROBiosystems) on HisPur™ Ni-NTA Magnetic Beads (Cat. # 88831, Thermo Fisher Scientific). The selection buffer consisted of 1x PBS, 0.05% Tween-20, 0.1 mg mL⁻¹ sssDNA, and 10 mM imidazole. Following two rounds of affinity selection, the eluted samples were submitted to WuXi AppTec for next-generation sequencing (NGS) and decoding. Hit identification was based on enrichment score (>1000) and copy number (>10). Data analysis identified DELC2 as a top-binding compound, whose tag-free version was synthesized by WuXi AppTec for subsequent investigation.

### Microscale Thermophoresis (MST)

Microscale thermophoresis (MST) was performed to evaluate direct binding between DELC2 (W1) and recombinant human CHI3L1. Recombinant His-tagged human CHI3L1 (Cat. #CH1-H5228, Acro Biosystems) was fluorescently labeled using the RED-tris-NTA 2nd Generation labeling kit (Cat. #MO-L018, NanoTemper Technologies) according to the manufacturer’s instructions. Labeled CHI3L1 was diluted to a final concentration of 80 nM in MST assay buffer containing 10 mM HEPES, 150 mM NaCl, 0.1% Pluronic F-127, 1 mM TCEP, and 4% DMSO (pH 7.4). DELC2 was prepared as a 16-point serial dilution ranging from 500 μM to the low nanomolar range and incubated with labeled protein for 30 min at room temperature. Following brief centrifugation, samples were loaded into premium capillaries and analyzed using a Monolith NT.115 instrument (NanoTemper Technologies). Thermophoretic measurements were collected under standard instrument settings, and binding affinities were determined using MO.Affinity Analysis v2.3 software. Concentration-dependent thermophoretic shifts were observed for DELC2, confirming direct interaction with CHI3L1 and yielding dissociation constants (K_D) in the low micromolar range.

### Surface Plasmon Resonance (SPR)

Surface plasmon resonance (SPR) analysis was performed using a Biacore™ 8K system (Cytiva, Marlborough, MA, USA) to validate direct binding between DELC2 and human CHI3L1. Recombinant His-tagged human CHI3L1 protein (8.0 μg mL^−1^) prepared in acetate buffer (pH 5.0) was immobilized onto a Series S Sensor Chip CM5 (Cytiva) using a standard amine-coupling protocol with a commercial amine coupling kit (BR100050, Cytiva). Immobilization was carried out at a flow rate of 10 μL min^−1^ for 420 s, yielding an immobilization level of approximately 3000 response units (RU), followed by blocking with ethanolamine. A reference flow cell treated with ethanolamine alone was used for background subtraction. PBS-P+ buffer (Cytiva) was used as the immobilization and running buffer (31).

Serial dilutions of DELC2 were prepared in assay buffer and injected over the immobilized CHI3L1 surface using a single-cycle kinetics protocol at a flow rate of 30 μL min^−1^ with an association time of 120 s per injection. Regeneration was performed for 30 s at 30 μL min^−1^ using regeneration buffer. Sensorgrams were processed and analyzed using Biacore™ Insight Evaluation Software (Cytiva). Binding experiments demonstrated concentration-dependent association and dissociation responses consistent with specific interaction between DELC2 and CHI3L1. Dissociation constants (K_D) were calculated from fitted binding curves and are reported as mean ± SD from independent measurements where indicated._

### Protein Expression and Purification of Recombinant Human CHI3L1

Human CHI3L1 (residues 1–383) containing a C-terminal HRV 3C protease cleavage site followed by a His-tag was cloned into a pCMV expression vector and transiently expressed in Expi293F™ suspension cells using the Expi293™ Expression System (Thermo Fisher Scientific). Expi293F™ cells were maintained in Expi293™ Expression Medium at 37 °C with 8% CO_2_ under orbital shaking conditions according to the manufacturer’s recommendations. Transient transfection was performed using the ExpiFectamine™ 293 Transfection Kit following the manufacturer’s protocol with a plasmid DNA-to-transfection reagent ratio of 1:5. Cells were cultured for 5 days post-transfection to allow recombinant protein expression prior to harvest.

Cell culture supernatants were collected by centrifugation and clarified prior to purification. Recombinant CHI3L1 was purified using Ni-NTA affinity chromatography with manually packed Ni-NTA resin. The clarified supernatant was incubated with Ni-NTA beads equilibrated in binding buffer containing 20 mM Tris-HCl (pH 8.0), 150 mM NaCl, 5% glycerol, and 10 mM imidazole. After extensive washing with binding buffer, bound protein was eluted using elution buffer supplemented with 300 mM imidazole. Eluted fractions containing CHI3L1 were pooled and further purified by size-exclusion chromatography (SEC) using a buffer containing 20 mM Tris-HCl (pH 8.0), 150 mM NaCl, and 5% glycerol. Purified protein fractions were concentrated to approximately 5 mg mL⁻¹ and subsequently used for crystallization experiments.

### Protein Crystallization, Data Collection, and Structure Determination

Purified recombinant human CHI3L1 was concentrated to 5 mg/mL in buffer containing 20 mM Tris-HCl (pH 8.0), 150 mM NaCl, and 5% glycerol prior to crystallization screening. Crystals of the CHI3L1-DELC2 complex were obtained using the sitting-drop vapor diffusion method at 289 K. Crystallization drops were prepared by mixing protein solution with reservoir solution containing 0.1 M HEPES (pH 7.0) and 10% (w/v) PEG 6000. Crystals appeared under conditions spanning a pH range of 5-7 and were subsequently optimized for diffraction studies.

X-ray diffraction data were collected at 100 K at the NSLS-II beamline 17-ID-1 using a Dectris EIGER X 9M detector and an X-ray wavelength of 0.97934 Å. Diffraction images were processed and scaled using autoPROC and Aimless (32, 33). The structure was solved by molecular replacement in PHASER using the previously reported CHI3L1 structure (PDB ID: 8R41) as the search model. Iterative model building and refinement were performed using PHENIX (version 1.21.1) ^33^. Structural analysis and visualization were carried out using standard crystallographic software packages. Data collection and refinement statistics are summarized in Table S1.

### CHI3L1-RAGE competition ELISA

High-binding 96-well plates were coated overnight at 4°C with recombinant human RAGE-Fc (2 μg/mL; 100 μL/well) in carbonate-bicarbonate buffer (50 mM, pH 9.6). Plates were washed three times with phosphate-buffered saline containing 0.05% Tween-20 (PBST) and blocked for 1 h at room temperature with PBST containing 3% bovine serum albumin. Recombinant human CHI3L1 (100 nM final concentration) was preincubated with DELC2 (0.01-100 μM) for 30 min at room temperature in assay buffer containing PBS, 0.05% Tween-20, 0.1% bovine serum albumin, and 1% DMSO. CHI3L1-DELC2 mixtures were transferred to RAGE-coated wells and incubated for 1 h at room temperature. After washing, retained CHI3L1 was detected using a rabbit anti-human CHI3L1 antibody followed by horseradish-peroxidase-conjugated anti-rabbit secondary antibody. Tetramethylbenzidine substrate was added for 10 min, and the reaction was stopped with 1 M sulfuric acid. Absorbance was measured at 450 nm with 570 nm reference correction. Background-subtracted values were normalized to the vehicle condition, defined as 100% CHI3L1-RAGE binding. Three independent experiments were performed at each DELC2 concentration. Apparent IC_50_ values were derived by four-parameter logistic regression in GraphPad Prism.

### Biolayer interferometry competition assay

BLI experiments were performed on an Octet R8 instrument using protein A biosensors. Biosensors were hydrated for at least 10 min in kinetics buffer consisting of PBS, 0.02% Tween-20, 0.1% bovine serum albumin, and 1% DMSO. Recombinant human RAGE-Fc (R&D Systems, Catalog # 1145-RG) was captured on biosensors at 10 μg/mL for 300 s, followed by a 60-s baseline step in kinetics buffer.

Recombinant human CHI3L1 (100 nM; ACROBiosystems, catalog no. CH1-H5228) was preincubated for 30 min at room temperature with vehicle or DELC2 at 0.1, 0.3, 1, 3, 10, or 30 μM. Biosensors were exposed to the mixtures for 120 s and then transferred to compound-matched kinetics buffer for 180 s. Reference biosensors without captured RAGE-Fc and buffer-only reference wells were included for double-reference subtraction. The response at the end of the 120-s association period was normalized to the vehicle response. Three independent experiments were performed at each concentration. Apparent IC_50_ values were calculated by four-parameter logistic regression.

### Astrocyte-microglia co-culture and CHI3L1-RAGE epistasis studies

Primary murine astrocytes and microglia were isolated from postnatal day 1-3 C57BL/6J mouse cortices. Cortices were dissociated enzymatically with papain and DNase I, triturated mechanically, and cultured in poly-D-lysine-coated T75 flasks in DMEM/F12 supplemented with 10% fetal bovine serum, 1% penicillin-streptomycin, and 1% GlutaMAX. Mixed glial cultures were maintained for 10-14 days. Microglia were collected by gentle orbital shaking, and adherent astrocytes were detached with trypsin-EDTA. Cells were plated in direct astrocyte-microglia co-culture at a 3:1 astrocyte-to-microglia ratio at a total density of 8 × 10^4^ cells per well in poly-D-lysine-coated 96-well plates. Independent cultures were prepared from three separate litters, with each litter-derived culture considered an independent biological replicate.

For RAGE epistasis experiments, cultures were pretreated for 30 min with DELC2 (10 μM), anti-RAGE neutralizing antibody (10 μg/mL), or matched isotype control before addition of recombinant murine CHI3L1 (500 ng/mL). A goat anti-mouse RAGE/AGER neutralizing antibody (R&D Systems, AF1179) or normal goat IgG isotype control (R&D Systems, AB-108-C) was used at 10 μg/mL. DELC2 was dissolved in DMSO; the final DMSO concentration was maintained at 0.1% in all treatment and vehicle-control wells. For NF-κB studies, cells were collected at 0, 15, 30, and 60 min. Phosphorylated NF-κB p65 (Ser536) and total NF-κB p65 were quantified in matched cell lysates using an AlphaLISA SureFire Ultra multiplex phospho-NF-κB p65 (Ser536)/total NF-κB p65 assay (Revvity). Cells were lysed directly in assay plates, and lysates were incubated with AlphaLISA detection reagents according to the manufacturer’s instructions. Time-resolved luminescence was measured on an EnVision multimode plate reader. Phospho-p65 signal was normalized to total p65 signal and expressed relative to vehicle-treated control cultures.

For delayed STAT3 studies, cells were collected at 0, 120, 240, and 480 min, and pSTAT3 (Tyr705) was quantified relative to total STAT3. Phosphorylated STAT3 (Tyr705) and total STAT3 were measured in matched lysates using an AlphaLISA SureFire Ultra multiplex phospho-STAT3 (Tyr705)/total STAT3 assay (Revvity). Luminescent phospho-STAT3/total-STAT3 ratios were normalized to the vehicle control. Culture supernatants collected 4 h after CHI3L1 stimulation were analyzed for IL-6 using a mouse IL-6 ELISA kit (R&D Systems, M6000B).

For assessment of microglial state and Aβ uptake, astrocyte-microglia co-cultures were incubated with HiLyte Fluor 488-labeled Aβ_42_ (500 nM) for 2 h at 37°C. Cells were washed extensively with ice-cold PBS to remove extracellular peptide, detached using Accutase, and stained with Zombie NIR Fixable Viability Dye, anti-CD11b, and anti-CD45. Cells were then fixed, permeabilized, and stained intracellularly with anti-Galectin-3 antibody. Samples were acquired on a BD LSRFortessa X-20 flow cytometer and analyzed using FlowJo software. Live singlet CD11b-positive, CD45-intermediate cells were defined as microglia. Galectin-3 positivity was established using fluorescence-minus-one controls. Aβ uptake was quantified as the median fluorescence intensity of HiLyte Fluor 488-positive microglia and expressed relative to vehicle-treated cultures.

### CHI3L1 neutralization control

For pathway-control experiments, astrocyte-microglia co-cultures were exposed to Aβ42 and recombinant CHI3L1 as described above and treated with an anti-CHI3L1 neutralizing antibody (10 μg/mL), low-dose DELC2 (1 μM), high-dose DELC2 (10 μM), or vehicle. A matched isotype antibody was included as control. A goat anti-mouse CHI3L1-neutralizing antibody (R&D Systems, AF2649) or matched normal goat IgG isotype control (R&D Systems, AB-108-C) was added 30 min before CHI3L1 stimulation. DELC2 and vehicle conditions contained 0.1% DMSO. Each condition was assessed in triplicate wells across three independently prepared primary glial cultures. Aβ uptake, DQ-BSA proteolysis, NF-κB reporter activity, and IL-1β secretion were quantified as described above.

### Computational study

Blind rigid-body docking was performed to assess whether the modeled RAGE V-C1 domain interface could spatially overlap the crystallographically defined DELC2-binding groove of CHI3L1. Ligand-free human CHI3L1 coordinates were derived from the CHI3L1-DELC2 crystal structure (PDB: 10KA; chain A; residues 22-383) after removal of DELC2. The human RAGE V-C1 domain was obtained from PDB: 3CJJ (chain A; residues 22-240). The DELC2 pocket was defined as the 20 CHI3L1 residues located within 4.5 Å of DELC2 in the 10KA complex.

Docking was conducted with LightDock version 0.9.4 using the DFIRE statistical potential without interface restraints or target-specific experimental constraints. Ten independent glowworm-swarm optimization searches, each initiated from a distinct surface position, were performed with 50 glowworms per swarm and 20 optimization steps per swarm. The calculations therefore sampled 500 starting poses. Docking was performed as a rigid-body analysis without anisotropic network model flexibility. The top-ranked pose from each swarm was retained for interface analysis. For the prespecified favorable-scoring subset, defined as swarms with positive DFIRE scores, CHI3L1 residues lying within 5 Å of RAGE in the top pose were identified and intersected with the crystallographic DELC2-pocket residue set. Five of ten swarms met the favorable-score criterion. Pocket overlap was observed in all five, and Trp99, Leu140, Tyr141, Asp207, Thr288, and Glu290 were shared across the four highest-scoring favorable swarms. Structural rendering and interface inspection were performed in PyMOL. This analysis was designed to generate a structural hypothesis and was not used to define the native CHI3L1-RAGE complex. Its interpretation is limited by blind, rigid-body sampling, use of a generic scoring potential, short optimization trajectories, analysis of one top pose per swarm, and incomplete convergence across all swarms.

### Human CSF studies

Commercially sourced de-identified cerebrospinal-fluid samples from donors with AD and non-demented controls were aliquoted on receipt and stored at -80°C. Samples underwent a single thaw cycle before use. CSF samples were obtained from PrecisionMed and included 12 donors with clinically diagnosed AD and 10 age-matched non-demented control donors. Samples were de-identified before purchase; their use was determined to be exempt from human-subjects review. CHI3L1 concentrations were quantified using a human CHI3L1 ELISA according to the manufacturer’s instructions using the Human Chitinase 3-like 1 Quantikine ELISA Kit (R&D Systems, DC3L10).

For functional studies, CSF was diluted to 10% (vol/vol) in serum-free glial culture medium. Astrocytes were exposed to individual control or Alzheimer’s disease CSF samples for 24 h. NF-κB reporter activity and fluorescent Aβ_42_ uptake were subsequently measured. Primary astrocytes were plated at 5 × 10^4^ cells per well in 96-well plates. DELC2 was added at 10 μM, and the final DMSO concentration was maintained at 0.1% across all conditions. For paired depletion studies, individual Alzheimer’s disease CSF aliquots were incubated overnight at 4°C with anti-CHI3L1 antibody-conjugated magnetic beads or isotype control antibody-conjugated beads. Beads were removed magnetically, and CHI3L1 depletion was confirmed by ELISA. Anti-human CHI3L1 antibody (R&D Systems, AF2599) or normal goat IgG control was coupled to Dynabeads Protein G (Thermo Fisher Scientific, 10004D) according to the manufacturer’s instructions. Depletion efficiency was calculated relative to the CHI3L1 concentration measured in the matched input aliquot. Recombinant human CHI3L1 was then added back to depleted CSF at 300 ng/mL. Matched aliquots from each donor were tested across untreated, DELC2-treated, depleted, and add-back conditions. For relay studies, astrocytes were exposed to CSF for 24 h, washed, and cultured for an additional 6 h in fresh medium. Conditioned medium was transferred to primary microglia for 18 h, after which IL-1β was measured by ELISA using the Mouse IL-1β Quantikine ELISA Kit (R&D Systems, MLB00C). Individual donors were treated as independent biological replicates.

### Oral pharmacokinetics and brain free fraction

All procedures were approved by the Institutional Animal Care and Use Committee of Weill Cornell Medicine (IACUC, Protocol #2024-0006). DELC2 was formulated as a suspension in 0.5% methylcellulose containing 0.1% Tween-80 and administered by oral gavage at 10 mg/kg. Male and female C57BL/6J mice were sampled at 0.25, 0.5, 1, 2, 4, 8, 12, and 24 h after dosing. Blood was collected by submandibular bleed or terminal cardiac puncture into K2-EDTA tubes and centrifuged at 2,000 × g for 10 min at 4°C to isolate plasma. Whole brains were harvested following transcardial perfusion with ice-cold PBS, weighed, snap-frozen, and stored at -80°C.

Brains were homogenized at 1:4 (wt/vol) in phosphate-buffered saline. DELC2 concentrations in plasma and brain homogenate were quantified by LC-MS/MS using matrix-matched calibration standards, quality-control samples, and an appropriate stable-isotope or structural analog internal standard. Chromatographic separation was performed using an Agilent 1290 Infinity II UHPLC coupled to an Agilent 6470 triple-quadrupole mass spectrometer and a Waters Acquity UPLC BEH C18 column (2.1 × 50 mm, 1.7 μm). Mobile phase A consisted of water with 0.1% formic acid, and mobile phase B consisted of acetonitrile with 0.1% formic acid. DELC2 was quantified in positive-ion multiple-reaction-monitoring mode using matrix-matched calibration curves spanning 1-10,000 ng/mL. Brain concentrations were corrected for homogenization dilution. Noncompartmental pharmacokinetic analyses were conducted using Phoenix WinNonlin version 8.3. The unbound fraction of DELC2 in mouse brain was determined by equilibrium dialysis. Brain homogenate was diluted fourfold in PBS and loaded into 96-well equilibrium-dialysis chambers with a 12-14 kDa molecular-weight-cutoff membrane. Equilibrium dialysis was conducted in the HTDialysis 96-well system using 150 μL diluted brain homogenate and 150 μL PBS receiver buffer per chamber. Samples were analyzed in triplicate. Homogenate and PBS receiver buffer were equilibrated for 6 h at 37°C with orbital shaking. DELC2 concentrations in donor and receiver compartments were measured by LC-MS/MS. The unbound fraction in homogenate was calculated as the receiver-to-donor concentration ratio and corrected for homogenate dilution to estimate the unbound fraction in intact brain. Unbound brain DELC2 concentrations were calculated by multiplying total brain concentration by the corrected free fraction.

### In vivo efficacy study using 5xFAD mice

All procedures were approved by the Institutional Animal Care and Use Committee of Weill Cornell Medicine (IACUC, Protocol #2024-0006). 5xFAD mice were obtained from The Jackson Laboratory (stock no. 034840), and transgene status was confirmed by PCR genotyping. Five-month-old male and female 5xFAD mice and age-matched wild-type littermate controls on a C57BL/6J background were assigned to WT vehicle, 5xFAD vehicle, 5xFAD + low-dose DELC2 (10 mg/kg), or 5xFAD + DELC2 (30 mg/kg) groups. DELC2 was administered once daily by oral gavage in 0.5% methylcellulose/0.1% Tween-80 for 21 consecutive days. Wild-type and 5xFAD vehicle groups received matched vehicle. Animals were balanced by sex across treatment groups. Investigators performing behavioral testing, tissue processing, and endpoint analysis were blinded to treatment allocation.

Y-maze testing was performed during the final treatment week. Mice were placed in the center of a three-arm maze and allowed to explore freely for 8 min. The maze contained three identical arms measuring 35 cm long × 5 cm wide × 15 cm high and was maintained at approximately 30 lux. Arm entries were recorded using EthoVision XT and were defined as placement of all four paws within an arm. Spontaneous alternation was calculated as the percentage of consecutive triads containing entries into all three arms relative to the maximum possible alternations. Total arm entries were recorded as an exploratory-activity control. Novel-object recognition was performed over three days. Mice were habituated to the testing arena on day 1, exposed to two identical objects on day 2, and tested 24 h later with one familiar and one novel object. The discrimination index was calculated as 100 × time exploring the novel object divided by total object-exploration time. Objects and arena positions were counterbalanced between animals, and the arena was cleaned with 70% ethanol between trials. Exploration was defined as directing the nose toward or touching an object within 2 cm; climbing or sitting on an object was not scored. Animals with less than 10 s total object-exploration time during testing were excluded according to the prespecified criterion. After behavioral testing, mice were euthanized and perfused with cold PBS. Cortex and hippocampus were dissected, snap-frozen, and stored at -80°C until biochemical analysis.

### Brain Aβ biochemical analyses

Cortical tissue was sequentially extracted in Tris-buffered saline (TBS; 50 mM Tris-HCl, 150 mM NaCl, pH 7.4) supplemented with protease and phosphatase inhibitor cocktails to obtain the TBS-soluble fraction, followed by 5 M guanidine hydrochloride extraction of the remaining pellet to obtain the insoluble fraction. Frozen cortex or hippocampus was homogenized at 100 mg tissue per mL of ice-cold extraction buffer using a bead-mill homogenizer. Homogenates were centrifuged at 100,000 × g for 30 min at 4°C to obtain the TBS-soluble fraction. Pellets were extracted in 5 M guanidine hydrochloride for 3 h at room temperature with end-over-end rotation and centrifuged at 20,000 × g for 20 min before collection of guanidine-soluble material. Aβ_42_ concentrations were quantified using the Invitrogen Amyloid beta 42 ELISA Kit (KHB3441), and Aβ_40_ concentrations were quantified using the Invitrogen Amyloid beta 40 ELISA Kit (KHB3481), according to the manufacturers’ instructions. Guanidine extracts were diluted to fall within the assay range before analysis. Aβ concentrations were normalized to total protein, determined by bicinchoninic acid assay (Pierce BCA Protein Assay Kit, Thermo Fisher Scientific, 23225), and reported as pg/mg protein. Total protein was measured separately in the matched soluble and guanidine-soluble fractions. Hippocampal Aβ_42_ and Aβ_40_ concentrations were quantified from matched extracts, and the Aβ_42_/Aβ_40_ ratio was calculated for each animal. Aggregated/oligomeric Aβ was quantified using the Invitrogen Human Amyloid beta (Aggregated) ELISA Kit (KHB3491) and expressed relative to the mean value of the WT-vehicle group.

### Bulk-brain CHI3L1 and glial inflammatory measures

Brain endogenous murine Chi3l1 expression was measured in TBS-soluble cortical homogenates by ELISA and expressed relative to the WT-vehicle mean. Frozen cortical tissue was homogenized at 100 mg/mL in ice-cold TBS containing protease and phosphatase inhibitors, centrifuged at 100,000 × g for 30 min at 4°C, and the resulting supernatants were aliquoted for all bulk-brain protein measurements. CHI3L1 was quantified using the Mouse Chitinase 3-like 1/YKL-40 Quantikine ELISA Kit (R&D Systems, MC3L10). Samples were diluted 1:10, analyzed in duplicate, normalized to total protein measured by Pierce BCA assay, and reported as pg/mg protein. Values below the lower limit of quantification were assigned the lower-limit value before analysis. Reactive astrocyte activity was summarized as a GFAP/C3 composite z score. GFAP was quantified using the Mouse/Rat GFAP Quantikine ELISA Kit (R&D Systems, MGF00), and C3 was quantified using a mouse Complement C3 ELISA kit (Abcam, ab263887). Matched cortical-extract aliquots were analyzed in duplicate, normalized to total protein, and converted to z scores using the mean and standard deviation of the WT-vehicle group. The GFAP/C3 composite score was calculated as the arithmetic mean of the two standardized values.

The pro-inflammatory microglial measure was quantified as the percentage of CD86-positive microglia in dissociated brain-cell preparations. An astrocyte inflammatory composite score was calculated from prespecified inflammatory markers measured in cortical lysates. This prespecified composite comprised CHI3L1, GFAP, C3, Serpina3n, and lipocalin-2. Serpina3n and lipocalin-2 were quantified using colorimetric mouse ELISAs in matched cortical-extract aliquots. Each normalized marker was converted to a z score relative to the WT-vehicle mean and standard deviation, and the final astrocyte inflammatory composite was calculated as the unweighted mean of the five z scores. Higher values indicate a more reactive astrocyte-associated inflammatory state.

### Brain dissociation and flow cytometry

Freshly collected brains were mechanically dissociated and enzymatically digested using papain and DNase I at 37°C for 30 min, followed by gentle trituration through fire-polished pipettes. Papain (Worthington Biochemical, LS003126) was used at 20 U/mL and DNase I (Sigma-Aldrich, DN25) at 50 U/mL. Cell suspensions were passed through 70-μm cell strainers and subjected to myelin removal using a density-gradient separation step. Myelin was removed using Myelin Removal Beads II (Miltenyi Biotec, 130-096-733). Cells were washed in ice-cold staining buffer consisting of PBS, 2% fetal bovine serum, and 2 mM EDTA.

After Fc receptor blocking, cells were stained with a fixable viability dye and antibodies against CD11b, CD45, TMEM119, P2RY12, ACSA-2, GLAST, and AGER/RAGE. Fc receptors were blocked with anti-mouse CD16/CD32 antibody (BioLegend, clone 93), and viability was assessed using Zombie NIR Fixable Viability Dye (BioLegend). Antibodies included anti-CD11b (clone M1/70), anti-CD45 (clone 30-F11), anti-TMEM119 (clone V.7C7), anti-P2RY12 (clone S16007D), anti-ACSA-2 (clone IH3-18A3), anti-GLAST (clone ACSA-1), and anti-RAGE (clone 175410). For intracellular phospho-protein staining, cells were fixed with 4% paraformaldehyde, permeabilized with methanol-based phospho-flow buffer, and stained with antibodies against pSTAT3 (Tyr705), phospho-NF-κB p65 (Ser536), and Galectin-3. Samples were acquired on a multicolor flow cytometer and analyzed using FlowJo software. Samples were acquired using a BD LSRFortessa X-20 and analyzed in FlowJo version 10.8.1. Single-color compensation controls and fluorescence-minus-one controls were included in every experiment. Doublets, debris, and nonviable cells were excluded before population gating. Microglia were defined as live CD11b-positive, CD45-intermediate, TMEM119-positive, P2RY12-positive cells. Astrocytes were defined as live ACSA-2-positive, GLAST-positive, CD11b-negative cells. Fluorescence-minus-one controls were used to establish positivity thresholds. Median fluorescence intensity was used for pSTAT3, pNF-κB, and AGER/RAGE measurements; the percentage of positive cells was used for Galectin-3 and CD86 endpoints.

### FACS purification and RT-qPCR

Freshly dissociated brain cells were prepared as described above. Following live-cell and singlet gating, astrocytes were sorted as ACSA-2-positive, GLAST-positive, CD11b-negative cells, whereas microglia were sorted as CD11b-positive, CD45-intermediate, TMEM119-positive, P2RY12-positive cells. Cell populations were collected directly into lysis buffer. Sorting was performed on a Sony SH800S cell sorter using a 100-μm microfluidic sorting chip. Cells were collected directly into Buffer RLT Plus containing β-mercaptoethanol. Post-sort purity was assessed on a small aliquot and exceeded 90% for each population. Purity was confirmed by reanalysis using the same gating hierarchy used for sorting. RNA was isolated using the RNeasy Micro Kit, including on-column DNase digestion. First-strand cDNA was synthesized from 50-100 ng total RNA using a high-capacity reverse-transcription kit. Quantitative PCR was performed with SYBR

Green chemistry on a QuantStudio 6 Real-Time PCR System. PowerUp SYBR Green Master Mix (Applied Biosystems) was used in technical duplicate reactions under the following program: 95°C for 2 min, followed by 40 cycles of 95°C for 15 s and 60°C for 60 s. Melt-curve analysis verified amplification specificity. Expression of Chi3l1, Ager, Stat3, Socs3, Nfkbia, Lgals3, C3, Il1b, Tnf, Trem2, and Apoe was normalized to Gapdh and Hprt1. The geometric mean of Gapdh and Hprt1 was used as the endogenous reference. Relative expression was calculated using the 2^-ΔΔCt^ method. Within each purified cell population, values were normalized to the corresponding WT-vehicle mean before log2 transformation for heatmap visualization.

### Statistical analysis

Data are presented as mean ± SEM unless otherwise indicated. Individual symbols represent independent biological replicates, individual CSF donors, or individual animals, as specified in the figure legends.

For two-group comparisons, two-sided unpaired Student’s t tests were used. For comparisons among multiple groups, one-way ANOVA with Tukey’s multiple-comparison correction was used. CHI3L1 epistasis experiments were analyzed by two-way ANOVA in CHI3L1-containing conditions, with DELC2 and anti-RAGE as independent factors; the interaction term tested non-additive effects. Matched CSF depletion/add-back experiments were analyzed by repeated-measures one-way ANOVA with Tukey’s post hoc testing. Associations between continuous variables were assessed using two-sided Pearson correlation analysis. Concentration-response curves were fitted with a four-parameter logistic model. Statistical analysis was performed in GraphPad Prism version 10. A two-sided p value below 0.05 was considered statistically significant.

## Results

### Identification of DELC2 through DNA-Encoded Library Screening

A DEL screening campaign using the DELopen library (4.2 billion compounds) was conducted against the human CHI3L1 protein, as previously described (13). Briefly, the selection workflow involved exclusion of non-target control (NTC) signals (C4) from the target conditions (C1-C3), followed by a multi-tiered filtering pipeline to remove compounds with elevated promiscuity risk and nonspecific binders (Figure 1A). Specifically, compounds with a Target Frequent Hitter Index (FHI) greater than 1, indicating higher molecular promiscuity risk, and a NTC FHI greater than 1, indicating higher matrix binding risk, were excluded prior to downstream enrichment analysis.

DEL family 10059 was selected for further analysis, representing one of the top three most enriched families, with 101 compounds identified out of the total 1,264 CHI3L1 binders. Sequencing-based building-block (BB) enrichment analysis of family 10059 revealed a dominant chemotype defined by BB182 at position 4, which was conserved in 55 of 101 selected compounds. In contrast to family 10034 from the prior campaign described (13),no single building block emerged as dominant at the remaining three positions, indicating greater structural diversity within the active binders at those sites.

Final compound selection was guided by two criteria: enrichment score and fraction of sp^3^-hybridized carbons (Fsp^3^). DELC2 was identified as the top candidate, exhibiting an enrichment score of 141,857.3 and an Fsp^3^ value of 0.56 (Figure 1B, 1C). The high Fsp3 reflects a predominantly three-dimensional molecular architecture, a feature associated with improved selectivity and favorable physicochemical properties relative to flat, aromatic-rich compounds. The compound was subsequently synthesized as a tag-free version by WuXi AppTec (purity >95%) and characterized by NMR, LC-MS, mass spectrometry, HPLC purity, and chiral SFC (Fig. S1-S5) to confirm its identity and suitability for downstream evaluation of CHI3L1 binding and functional inhibition.

Microscale thermophoresis (MST) was used to assess direct binding of DELC2 to recombinant His-tagged human CHI3L1. DELC2 produced a concentration-dependent thermophoretic response, yielding a low-micromolar dissociation constant (K_D_ = 10.31 ± 6.4 μM; Figure 1D). To further validate the direct interaction between CHI3L1 and the top hit compound DELC2, surface plasmon resonance (SPR) analysis was performed under equilibrium binding conditions. Recombinant CHI3L1 was immobilized on an CM5 sensor chip, and serial dilutions of DELC2 were injected to monitor real-time association and dissociation kinetics. The resulting sensorgrams displayed a concentration-dependent increase in response units, consistent with specific and reversible target engagement (Figure 1E, 1F). Global fitting of the binding responses yielded an apparent dissociation constant (K_D_) of 8.7 ± 0.70 μM, indicating a high-affinity interaction for a small molecule targeting an extracellular glycan-recognition protein. Importantly, MST counterscreen data across 25 proteins suggested preferential DELC2 binding to CHI3L1, with weak or no quantifiable binding over a broad panel of related inflammatory, extracellular, and AD-relevant proteins (Figure S6).

**Figure 1.**
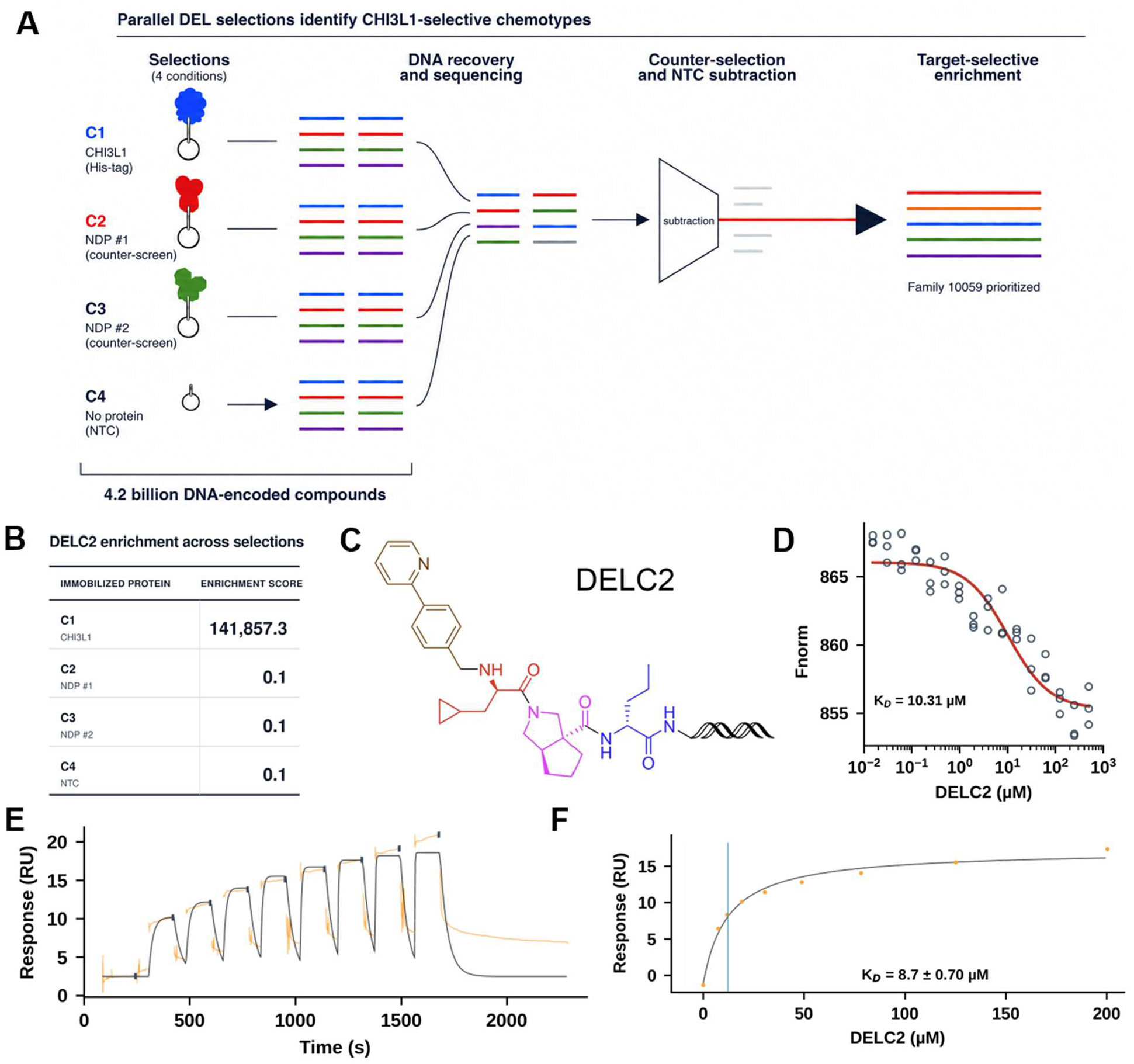
DNA-encoded library discovery and orthogonal biophysical validation of the CHI3L1-binding small molecule DELC2. **(A)** Twenty-seven DNA-encoded libraries comprising approximately 4.2 billion compounds were screened against recombinant human CHI3L1 immobilized on HisPur Ni-NTA magnetic beads (C1). Two immobilized non-disclosed human proteins (C2 and C3) served as counter-selections, and the affinity matrix alone served as the no-target control (C4). Following affinity selection, retained compounds were decoded by DNA-barcode sequencing. CHI3L1-enriched chemotypes were prioritized after counter-selection and no-target subtraction. **(B)** DELC2 enrichment across the four selection conditions. **(C)** Chemical structure of DELC2. **(D)** Microscale thermophoresis (MST) analysis of DELC2 binding to recombinant human CHI3L1. Nonlinear fitting of the concentration-dependent thermophoretic response yielded a binding affinity of 10.31 µM. **(E)** Representative SPR sensorgram for DELC2 binding to immobilized human CHI3L1. **(F)** Steady-state SPR analysis, yielding an apparent 8.7 µM. Together, MST and SPR provide orthogonal evidence of direct low-micromolar engagement of human CHI3L1 by DELC2. MST and SPR measurements were performed in three independent experiments.

### Structural basis of DELC2 recognition by CHI3L1

The crystal structure of CHI3L1 in complex with DELC2 (PDB ID: 10KA) revealed that DELC2 occupies the canonical glycan-recognition groove and stabilizes a compact aromatic-hydrophobic architecture within the ligand-binding cleft (Figure 2). Clear and continuous 2F_o_-F_c_ electron density contoured at 1σ enabled unambiguous placement of DELC2 within the elongated groove characteristic of chitinase-like proteins (Figure 2a,b). Data collection and refinement statistics are summarized in Table S1.

The distal region of DELC2 extends toward the solvent-accessible surface of the groove and engages residues surrounding the conserved Asp207/Tyr206 and Arg263 recognition interfaces, regions previously implicated in glycan and inhibitor binding (Figure 2c,ci). These interactions bridge the central hydrophobic cavity with the distal polar surface of the cleft, allowing DELC2 to span multiple recognition subsites simultaneously. Rather than relying on localized electrostatic anchoring alone, DELC2 appears to stabilize the groove through coordinated aromatic enclosure together with distributed polar complementarity across the cleft.

Rather than functioning as a classical carbohydrate mimic, DELC2 exploits the intrinsic hydrophobic topology of the CHI3L1 recognition cleft through extensive aromatic enclosure. The ligand is positioned within a densely packed aromatic environment centered on Trp99 and Trp352, two residues that constitute the principal anchoring elements of the glycan-recognition groove. The bicyclic core of DELC2 becomes embedded within an aromatic-hydrophobic cage formed by Trp99, Trp352, Phe261, Tyr141, Tyr206, Leu140, and adjacent hydrophobic residues, generating extensive π-stacking and shape-complementary packing interactions across the cleft (Figure 2c,cii). This mode of recognition differs substantially from previously described carbohydrate-mimetic CHI3L1 ligands and instead reveals a mechanism driven by broad hydrophobic occupancy and conformational stabilization of the groove architecture itself.

A defining feature of the structure is the extensive engagement of Trp99, a conformationally dynamic residue previously shown to undergo substantial rearrangement during carbohydrate accommodation. Earlier structural studies established Trp99 as a gatekeeper of the CHI3L1 recognition cleft, capable of adopting distinct conformational states during ligand binding. The rigid and conformationally constrained scaffold of DELC2 appears ideally suited to stabilize this region through extensive aromatic packing interactions, consistent with formation of a ligand-locked groove architecture. Together with the deep hydrophobic enclosure surrounding the bicyclic core, these observations support a model in which DELC2 suppresses the intrinsic conformational plasticity of the CHI3L1 carbohydrate-recognition cleft by stabilizing a compact Trp99/Trp352-centered recognition state.

Comparison with previously reported CHI3L1-carbohydrate and inhibitor complexes further emphasizes the distinct nature of DELC2 recognition. Whereas earlier CHI3L1 ligands primarily relied on carbohydrate mimicry and localized interaction hotspots, DELC2 achieves binding through extended aromatic occupancy and hydrophobic shape complementarity that effectively reorganize the dynamic recognition cleft. The ligand additionally engages regions proximal to the His209-Trp212 conformational axis implicated in ligand-induced cleft remodeling, suggesting that DELC2 binding may stabilize a groove conformation that is incompatible with normal carbohydrate-accommodation dynamics. Although the precise dynamic consequences of DELC2 engagement remain to be fully resolved, the 10KA structure identifies a ligand-induced aromatic-cage stabilization mechanism in CHI3L1 in which small-molecule binding constrains the conformational flexibility of the glycan-recognition groove.

The DELC2-bound structure defines the glycan-recognition groove of CHI3L1 as a druggable regulatory site and identifies a distributed aromatic, hydrophobic, and polar interaction network that anchors the ligand within this cleft. Although the structure does not by itself establish how DELC2 affects receptor engagement, occupancy of this ligand-accessible surface provides a structural rationale for testing whether DELC2 alters CHI3L1-dependent signaling through RAGE. We therefore next examined whether pharmacologic CHI3L1 targeting converges functionally with RAGE blockade in glial systems.

**Figure 2.**
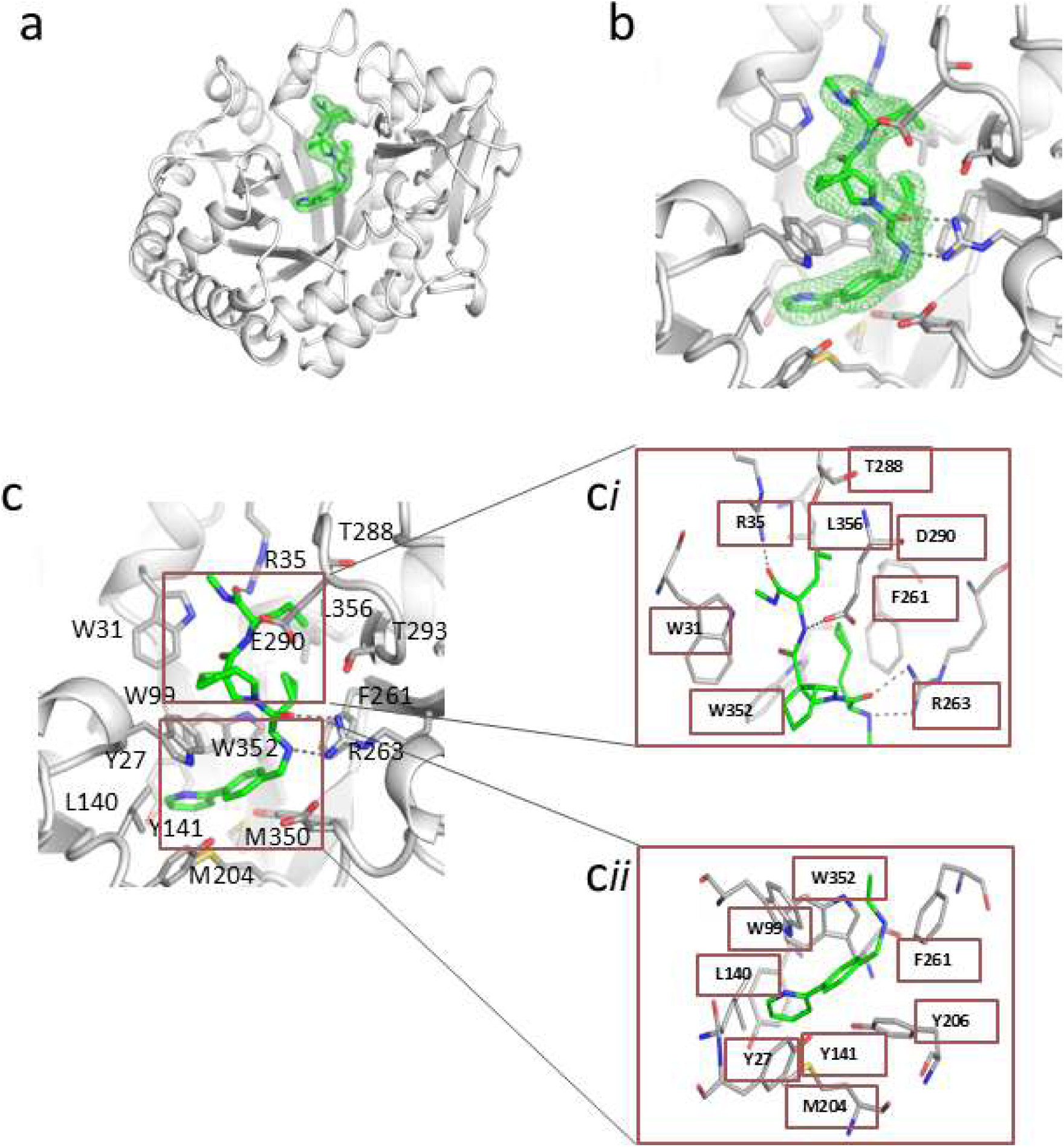
Structural basis of DELC2 recognition by CHI3L1. **(A)** Overall crystal structure of human CHI3L1 in complex with DELC2 (green), showing ligand occupancy within the canonical glycan-recognition groove. CHI3L1 is shown in cartoon representation. **(B)** Enlarged view of the DELC2-binding pocket with the 2F_o_-F_c_ electron density map contoured at 1σ, confirming unambiguous ligand placement within the cleft. **(C)** Detailed view of the DELC2 interaction network within the CHI3L1 recognition groove. DELC2 engages multiple residues spanning both polar and hydrophobic regions of the cleft. **(Ci)** Close-up of distal polar interactions involving residues Arg35, Thr288, Asp290, Leu356, Phe261, and Arg263. Dashed lines indicate potential hydrogen-bonding and polar contacts. **(Cii)** Close-up of the hydrophobic-aromatic enclosure surrounding DELC2 formed by Trp99, Trp352, Phe261, Tyr141, Tyr206, Leu140, and Met204, highlighting extensive aromatic packing interactions within the binding groove.

### DELC2 reduces CHI3L1-RAGE engagement and converges functionally with RAGE blockade to restrain glial inflammatory signaling

The DELC2-bound CHI3L1 structure identifies the glycan-recognition groove as the site of small molecule engagement (Figure 2). To determine whether occupancy of this groove affects receptor engagement, we measured CHI3L1-RAGE association in two orthogonal competition formats. In an ELISA using immobilized RAGE and fixed CHI3L1, DELC2 produced a concentration-dependent reduction in CHI3L1 retention, with an apparent competition IC_50_ of 2.25 ± 0.17 μM (Figure 3A). Consistently, biolayer interferometry (BLI) using a RAGE-Fc biosensor showed progressive attenuation of the CHI3L1-associated response with increasing DELC2 concentration (Figure 3B). Endpoint analysis at the end of the association phase yielded an apparent IC_50_ of 2.02 ± 0.19 μM (Figure 3C). These complementary assays support the conclusion that DELC2 reduces CHI3L1-RAGE engagement.

We next examined whether this biochemical effect is reflected in the downstream inflammatory response. Recombinant CHI3L1 induced an early increase in NF-κB p65 phosphorylation, which was attenuated by DELC2 and by RAGE blockade (Figure 3D). In CHI3L1-containing conditions, the DELC2 × anti-RAGE interaction was significant (two-way ANOVA, *p* = 0.0169), consistent with functional convergence at the early NF-κB response. CHI3L1 also increased IL-6 secretion at 4 h; both DELC2 and anti-RAGE reduced this response, with a significant DELC2 × anti-RAGE interaction (*p* < 0.0001; Figure 3E). CHI3L1-induced signaling extended to delayed STAT3 activation. DELC2 and anti-RAGE each attenuated pSTAT3 accumulation across the late time course (Figure 3F). However, the DELC2/anti-RAGE interaction did not reach significance at this downstream endpoint (*p* = 0.1467), indicating overlapping functional effects without a demonstrable interaction under these assay conditions. As an independent pathway control, direct CHI3L1 neutralization similarly reduced CHI3L1-associated inflammatory activity and restored Aβ handling in the astrocyte-microglia co-culture system (Figure S7).

To explore the structural basis for the observed competition, we performed blind docking of the RAGE V-C1 domain to ligand-free CHI3L1. Across favorable-scoring solutions, the modeled RAGE-contact region recurrently involved residues within the crystallographically defined DELC2-binding groove, including Trp99, Leu140, Tyr141, Asp207, Thr288, and Glu290 (Figure S8). This analysis is consistent with partial spatial overlap between the DELC2 pocket and a putative RAGE-contact surface. Because the docking was unrestrained and rigid-body, and interface convergence was incomplete across all swarms, it is interpreted as hypothesis-generating rather than as a structural definition of the CHI3L1-RAGE complex. Therefore, the combined results from the crystal structure, orthogonal competition assays, exploratory interface modeling, and RAGE epistasis experiments support a model in which DELC2 occupies the CHI3L1 glycan-recognition groove and reduces CHI3L1-RAGE engagement, thereby suppressing RAGE-linked NF-κB, IL-6, and STAT3 signaling.

**Figure 3.**
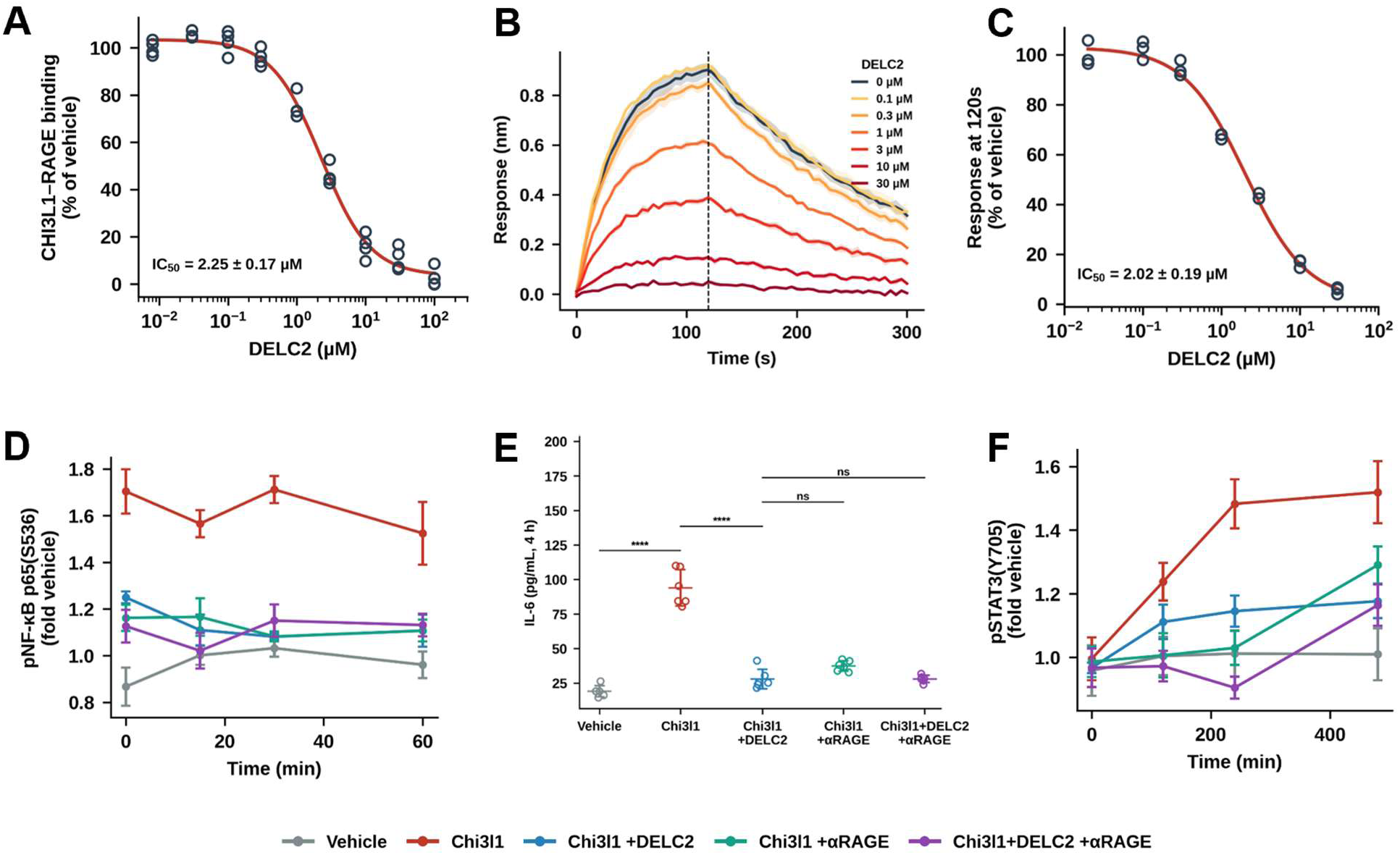
DELC2 reduces CHI3L1-RAGE engagement and suppresses RAGE-linked inflammatory signaling. **(A)** ELISA-based competition assay using immobilized RAGE and fixed recombinant CHI3L1. DELC2 reduced CHI3L1-RAGE binding in a concentration-dependent manner. **(B)** Representative BLI sensorgrams using a RAGE-Fc biosensor. Fixed CHI3L1 was preincubated with the indicated DELC2 concentrations before injection; the dashed line marks the end of the association phase. **(C)** BLI association-endpoint analysis at 120 s, showing a concentration-dependent decrease in the CHI3L1-associated response. **(D)** Time-resolved NF-κB p65 phosphorylation following CHI3L1 stimulation in the presence of vehicle, DELC2, anti-RAGE, or the combination. **(E)** IL-6 secretion measured 4 h after CHI3L1 stimulation. **(F)** Time-resolved STAT3 phosphorylation. For panels **D-F**, two-way ANOVA was performed in CHI3L1-containing arms with DELC2 and anti-RAGE as factors; interaction *p* values were 0.0169 for NF-κB, <0.0001 for IL-6, and 0.1467 for pSTAT3. For panels **A-C**, each concentration was tested in three independent experiments. For panels **D-F**, data are mean ± SEM from five independent biological replicates per condition. Statistical significance is denoted as follows: ns, not significant and ****p < 0.0001.

### DELC2 disrupts the CHI3L1-driven astrocyte-microglia inflammatory relay and restores Aβ handling

Because the CHI3L1-RAGE axis is positioned upstream of glial inflammatory signaling, we next examined whether DELC2 alters the downstream microglial state and its functional consequences for Aβ handling. In the astrocyte-microglia co-culture system, recombinant CHI3L1 increased the proportion of Gal-3-positive microglia relative to vehicle-treated cultures, consistent with induction of a pro-inflammatory microglial state (Figure 4A). DELC2 significantly reduced this CHI3L1-associated Gal-3 response. RAGE blockade produced a similar reduction, and combined DELC2 plus anti-RAGE treatment further reduced the Gal-3-positive population. The absence of a significant difference between DELC2 and anti-RAGE alone indicates that the two interventions exert broadly overlapping effects on this CHI3L1-dependent microglial response.

CHI3L1 exposure also impaired microglial Aβ uptake (Figure 4B). DELC2 restored Aβ uptake relative to the CHI3L1 condition, whereas RAGE blockade produced a comparable recovery. Addition of anti-RAGE to DELC2 did not produce a further statistically detectable improvement in uptake, consistent with convergence of DELC2 and RAGE blockade on a common CHI3L1-linked pathway. These results extend the biochemical competition and proximal signaling findings in Figure 3 to a functional glial outcome: DELC2 restrains the transition toward a Gal-3-associated inflammatory microglial state while restoring Aβ handling. Direct CHI3L1 neutralization served as an independent pathway control. In parallel astrocyte-microglia co-cultures, a CHI3L1-neutralizing antibody reduced NF-κB activity and IL-1β secretion, improved lysosomal proteolysis, and restored Aβ uptake, phenocopying the direction of DELC2 activity (Figure S7). Together with the CHI3L1-RAGE competition assays and RAGE epistasis analysis (Figure 3), these data support a model in which DELC2 occupancy of the CHI3L1 glycan-recognition groove reduces CHI3L1-RAGE engagement, thereby limiting propagation of an astrocyte-microglia inflammatory relay and preserving microglial Aβ handling.

**Figure 4.**
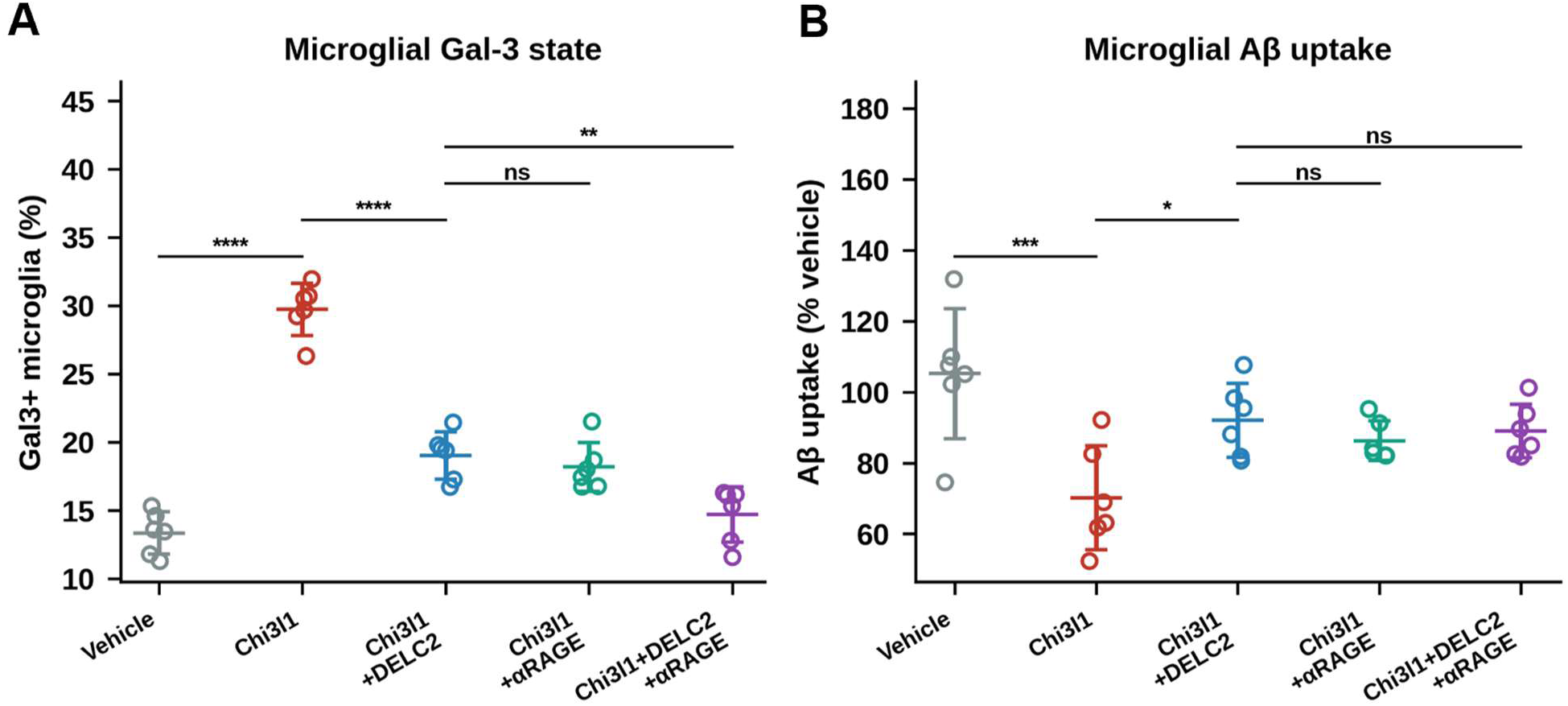
DELC2 restrains a CHI3L1-driven inflammatory microglial state and restores Aβ uptake. **(A)** Percentage of Gal-3-positive microglia in the astrocyte-microglia co-culture system, quantified by flow cytometry, after exposure to vehicle, recombinant CHI3L1, CHI3L1 plus DELC2, CHI3L1 plus anti-RAGE, or CHI3L1 plus the DELC2/anti-RAGE combination. **(B)** Microglial Aβ uptake under the same conditions, expressed relative to vehicle-treated cultures. Data are mean ± SEM from five independent biological replicates per condition; individual symbols represent independent replicates. Statistical significance is denoted as follows: ns, not significant; *p < 0.05; **p < 0.01; ***p < 0.001; and ****p < 0.0001.

### DELC2 suppresses CHI3L1-associated inflammatory activity in AD CSF

To examine the translational relevance of the CHI3L1-RAGE axis, we analyzed commercially sourced cerebrospinal fluid (CSF) samples from donors with AD and non-demented controls. CHI3L1 concentrations were significantly higher in AD CSF than in control CSF (Figure 5A). Across the donor cohort, CSF CHI3L1 concentration positively correlated with astrocyte NF-κB activity following CSF exposure (*r* = 0.62, *p* = 0.0012; Figure 5B), linking elevated CHI3L1 to a pro-inflammatory glial response.

We next tested whether DELC2 could attenuate the activity of AD CSF in glial cultures. Relative to control CSF, AD CSF increased astrocyte NF-κB activity (Figure 5C), reduced astrocytic Aβ uptake (Figure 5D), and increased microglial IL-1β secretion (Figure 5E). DELC2 attenuated each of these AD-CSF-associated phenotypes. Immunodepletion of CHI3L1 from matched AD-CSF aliquots similarly reduced NF-κB activation and microglial IL-1β secretion and restored astrocytic Aβ uptake. Reconstitution of CHI3L1-depleted AD CSF with recombinant CHI3L1 reinstated the inflammatory and Aβ-handling phenotypes. The paired depletion-add-back design establishes CHI3L1 as a functional contributor to the glial activity of AD CSF in this experimental system.

**Figure 5.**
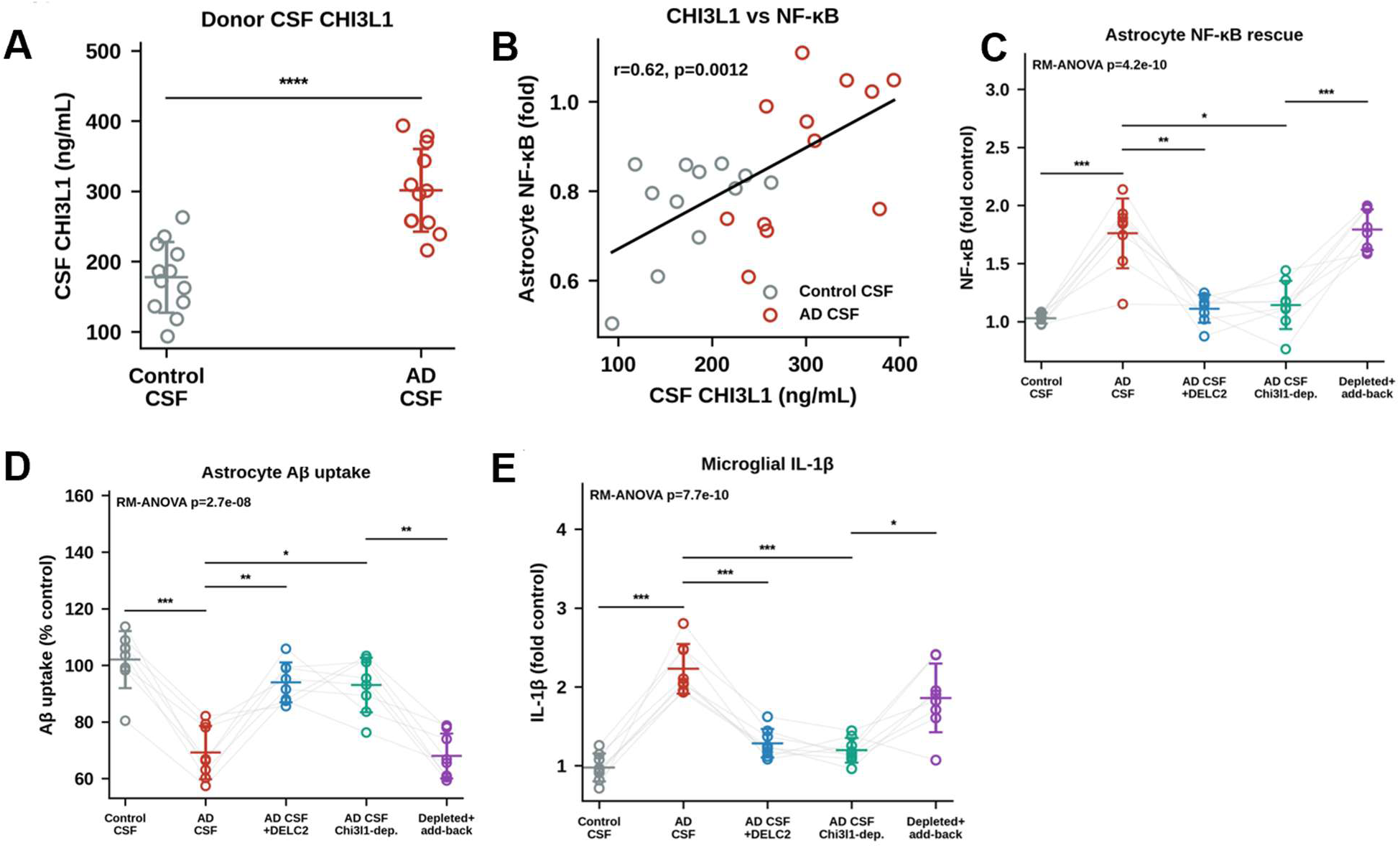
CHI3L1 is elevated in AD CSF and contributes to CSF-associated glial inflammatory activity. **(A)** CHI3L1 concentrations in commercially sourced CSF from non-demented control donors and donors with AD. **(B)** Correlation between CSF CHI3L1 concentration and astrocyte NF-κB activity after exposure to individual CSF samples. **(C-E)** Functional testing of matched AD-CSF aliquots after treatment with DELC2, CHI3L1 immunodepletion, or CHI3L1 depletion followed by recombinant CHI3L1 add-back. **(C)** Astrocyte NF-κB activity. **(D)** Astrocytic Aβ uptake. **(E)** Microglial IL-1β secretion. Gray lines connect matched aliquots from the same AD-CSF donor across conditions. For panels **A** and **B**, control CSF, n = 10 donors; AD CSF, n = 12 donors. For panels **C**-**E**, n = 12 matched AD-CSF donors per condition. Individual symbols represent independent donor samples; bars indicate mean ± SEM. Repeated-measures ANOVA and post hoc comparisons were performed as indicated. Statistical significance is denoted as follows: ns, not significant; *p < 0.05; **p < 0.01; ***p < 0.001; and ****p < 0.0001.

### Oral DELC2 administration produces sustained unbound brain exposure

We next characterized the pharmacokinetic profile of DELC2 after oral administration of the optimized formulation (10 mg/kg, once daily). DELC2 was rapidly detected in plasma and brain, with brain concentrations exceeding plasma concentrations throughout the 24-h sampling period (Figure 6A). Brain exposure was sustained, with measurable concentrations remaining at 24 h after dosing.

Measurement of the unbound fraction in brain homogenate yielded a mean brain free fraction (*f*_u,_brain) of 0.114 ± 0.024 (Figure 6B). Integration of individual brain-concentration profiles with the measured free fraction demonstrated sustained unbound brain DELC2 concentrations over the dosing interval (Figure 6C). Unbound brain exposure remained in the micromolar range at 24 h, providing pharmacokinetic support for once-daily dosing in subsequent in vivo studies. Consistent with its oral exposure profile, DELC2 displayed favorable in vitro ADME and early safety properties (Table S2). DELC2 showed high kinetic solubility in PBS (212 μM) and FaSSIF (318 μM), good passive permeability (PAMPA P_app_, 18.7 × 10^-6^ cm/s), stability in simulated gastric and intestinal fluids, and moderate-to-high microsomal stability in mouse and human liver microsomes (t_1/2_, 75.3 and 92.5 min, respectively). DELC2 was stable in mouse and human plasma and PBS over the 120-min assay window, showed 92.3% human plasma protein binding, and exhibited no measurable cytotoxicity in WI-38 or HS-27 cells or hERG inhibition at concentrations up to 100 μM. These data establish that oral administration of DELC2 achieves sustained brain exposure at a dose compatible with daily in vivo dosing. The exposure analysis supports pharmacological evaluation in 5xFAD mice but does not itself establish in vivo target occupancy.

**Figure 6.**
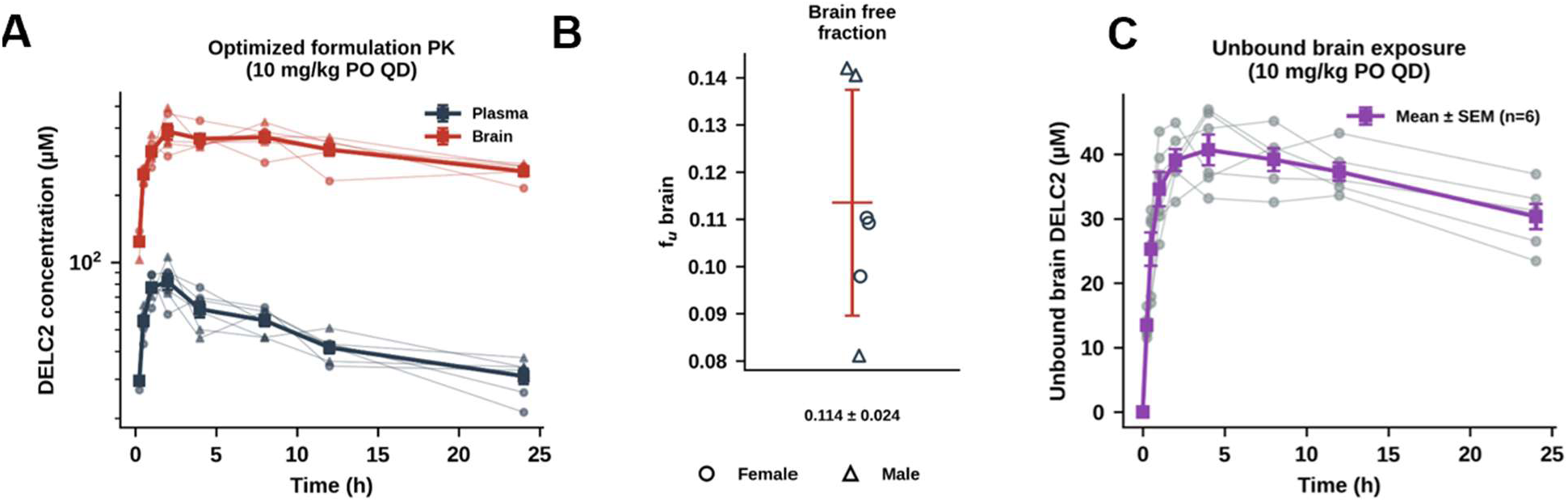
Oral pharmacokinetics and unbound brain exposure of DELC2. **(A)** Plasma and total brain concentration-time profiles after oral administration (n = 6 mice) of the DELC2 formulation (10 mg/kg). Thin lines show individual animals and symbols/lines show mean ± SEM. **(B)** Brain free fraction (*f*_u_,brain) determined in independent brain-homogenate equilibrium-dialysis measurements; circles and triangles indicate female and male animals (n = 6), respectively. Bars indicate mean ± SD. **(C)** Individual and mean ± SEM unbound brain DELC2 concentration-time profiles, calculated from total brain concentrations and the measured *f*_u_,brain.

### DELC2 improves cognitive and amyloid-associated outcomes while suppressing glial inflammatory programs in 5xFAD mice

Before efficacy studies, we confirmed that DELC2 engages recombinant murine Chi3l1 using MST (Figure S9). This cross-species target-engagement result provides a pharmacological bridge between the human CHI3L1 structural and competition studies and the 5xFAD model. Together with the sustained unbound brain exposure achieved after oral administration (Figure 6), these data supported in vivo evaluation of DELC2.

Relative to wild-type vehicle-treated controls, 5xFAD vehicle-treated mice displayed impaired spontaneous alternation in the Y-maze and reduced discrimination in the novel-object-recognition task (Figure 7A,B). DELC2 improved performance in both paradigms, with the higher dose producing the most consistent effect. Total arm entries during the Y-maze did not differ across groups (one-way ANOVA, p = 0.63), indicating that the behavioral effects were not attributable to altered exploratory activity. Concordant improvement across these complementary behavioral measures supports an effect on disease-relevant cognitive dysfunction rather than an isolated change in a single behavioral endpoint.

We next examined whether the behavioral effects were accompanied by changes in cerebral Aβ biochemistry. As expected, 5xFAD vehicle-treated mice exhibited elevated cortical soluble Aβ_42_, cortical insoluble Aβ_42_, hippocampal Aβ_42_/Aβ_40_ ratio, and oligomeric Aβ abundance relative to wild-type controls (Figure 7C-F). DELC2 reduced each of these amyloid-associated abnormalities, with the higher dose again producing the most consistent response across endpoints. The reduction in soluble Aβ_42_ and oligomeric Aβ is particularly notable because these species are implicated in synaptic dysfunction and inflammatory amplification. The accompanying reduction in insoluble Aβ_42_ and normalization of the hippocampal Aβ_42_/Aβ_40_ ratio indicate that sustained DELC2 treatment altered the biochemical balance of Aβ species in brain tissue. However, because quantitative plaque histology was not performed, these findings should not be interpreted as direct evidence of plaque clearance or remodeling. The in vivo biochemical effects were mechanistically coherent with the cellular findings. DELC2 reduced the CHI3L1-associated Gal-3-positive microglial state and restored microglial Aβ uptake in astrocyte-microglia co-cultures. It also attenuated AD-CSF-associated NF-κB activation and inflammatory cytokine production while improving astrocytic Aβ uptake. Thus, the reduction in soluble and oligomeric Aβ in 5xFAD brain is consistent with improved glial handling of Aβ and reduced inflammatory reinforcement, although the present experiments do not distinguish altered production, clearance, aggregation, or distribution of individual Aβ pools.

To determine whether these efficacy-associated outcomes were accompanied by suppression of glial inflammatory activation in brain, we quantified bulk-brain pharmacodynamic markers. Brain Chi3l1 abundance was elevated in 5xFAD vehicle-treated mice and was reduced following DELC2 treatment (Figure 7G). DELC2 also reduced the A1-reactive astrocyte composite score, the pro-inflammatory microglial measure, and the astrocyte inflammatory composite score (Figure 7H-J). The coordinated reduction in these markers is consistent with attenuation of a broader glial inflammatory state rather than modulation of a single downstream readout. These bulk-brain changes provide an in vivo correlate of the astrocyte-microglia relay identified in the co-culture system.

**Figure 7.**
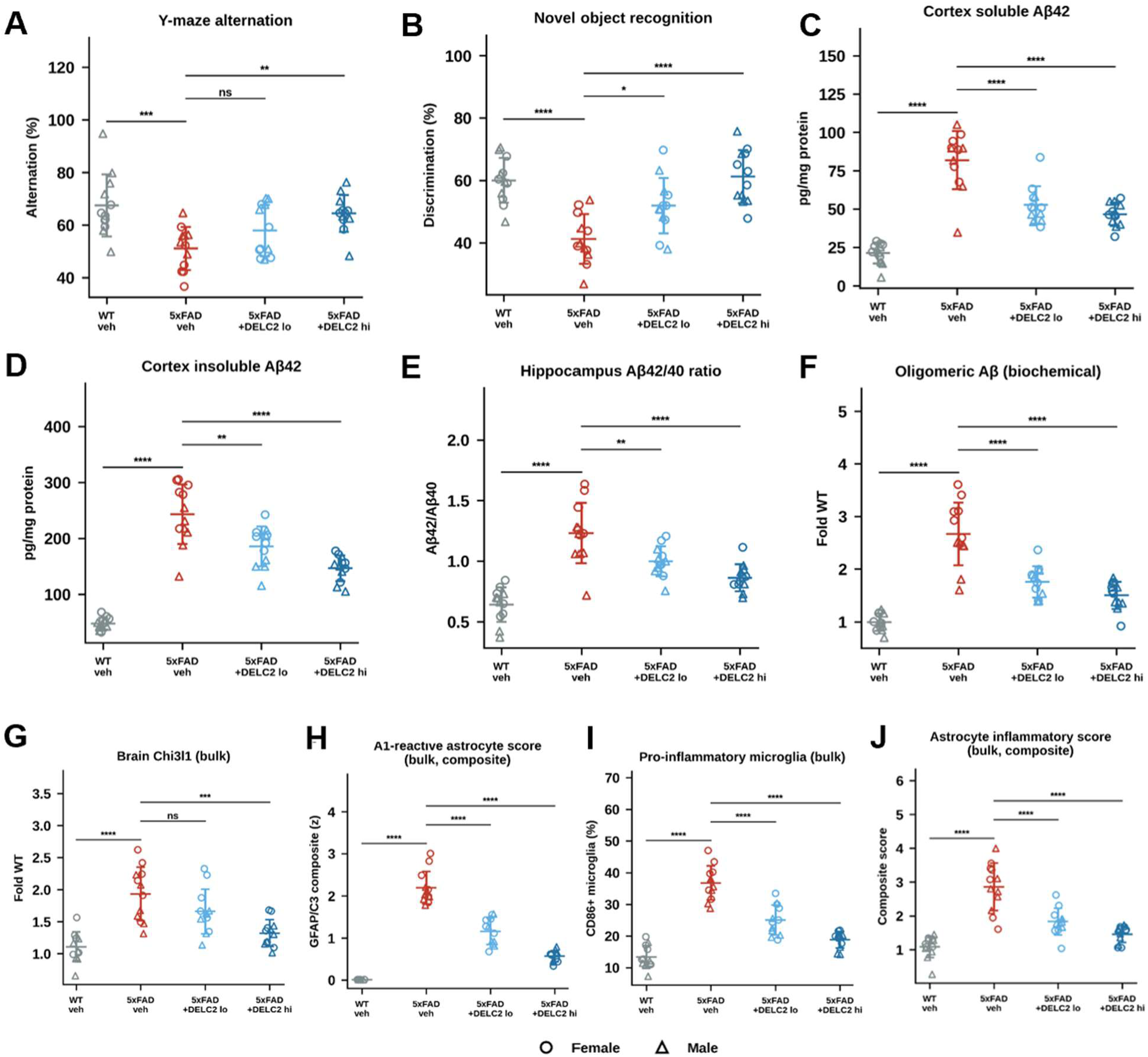
DELC2 improves cognitive and amyloid-associated outcomes while suppressing glial inflammatory programs in 5xFAD mice (n = 12 mice per group). **(A)** Y-maze spontaneous alternation. **(B)** Novel-object-recognition discrimination performance. **(C)** Cortical soluble Aβ_42_. **(D)** Cortical insoluble Aβ_42_. **(E)** Hippocampal Aβ_42_/Aβ_40_ ratio. **(F)** Oligomeric Aβ measured by biochemical assay. **(G)** Bulk-brain Chi3l1 abundance. **(H)** GFAP/C3 reactive-astrocyte composite score. **(I)** Percentage of CD86-positive microglia. **(J)** Astrocyte inflammatory composite score. Wild-type vehicle and 5xFAD vehicle groups were compared with 5xFAD mice treated with DELC2 at the indicated dose levels. Circles indicate female mice and triangles indicate male mice; each symbol represents one animal. Bars indicate mean ± SEM. Statistical significance is denoted as follows: ns, not significant; *p < 0.05; **p < 0.01; ***p < 0.001; and ****p < 0.0001.

### DELC2 normalizes cell-resolved astrocyte and microglial inflammatory signaling states in 5xFAD brain

Because the bulk-brain pharmacodynamic effects of DELC2 could not assign signaling changes to specific glial populations, we next analyzed dissociated brain cells by flow cytometry. Compared with WT vehicle controls, 5xFAD vehicle mice exhibited increased astrocytic pSTAT3 and pNF-κB mean fluorescence intensity (MFI), indicating activation of two central inflammatory signaling pathways in astrocytes (Figure 8A,B). DELC2 treatment reduced both readouts, with the high-dose group showing the most consistent normalization toward the WT range. These cell-resolved data extend the bulk-tissue findings and demonstrate that DELC2-associated suppression of inflammatory signaling includes astrocytes.

Microglial activation was similarly attenuated by DELC2. The proportion of Gal-3-positive microglia was increased in 5xFAD vehicle brains and was reduced by DELC2 treatment, with a greater effect at the high dose (Figure 8C). DELC2 also reduced the proportion of pSTAT3-positive microglia (Figure 8D). The coordinated reduction in Gal-3 and pSTAT3 supports attenuation of a disease-associated inflammatory microglial state and is consistent with the restoration of Aβ handling observed in the astrocyte-microglia co-culture system. Finally, Ager/RAGE signal in the analyzed brain-cell suspension was elevated in 5xFAD vehicle mice and reduced following DELC2 treatment (Figure 8E). In the context of the biochemical competition assays, RAGE epistasis experiments, and bulk-brain pharmacodynamic changes, this finding supports a model in which DELC2 reduces activity of a CHI3L1-RAGE-linked inflammatory axis in the 5xFAD brain. Although these flow-cytometry analyses do not establish direct target occupancy in individual cell populations, they demonstrate coordinated normalization of astrocytic and microglial inflammatory signaling states following oral DELC2 treatment.

**Figure 8.**
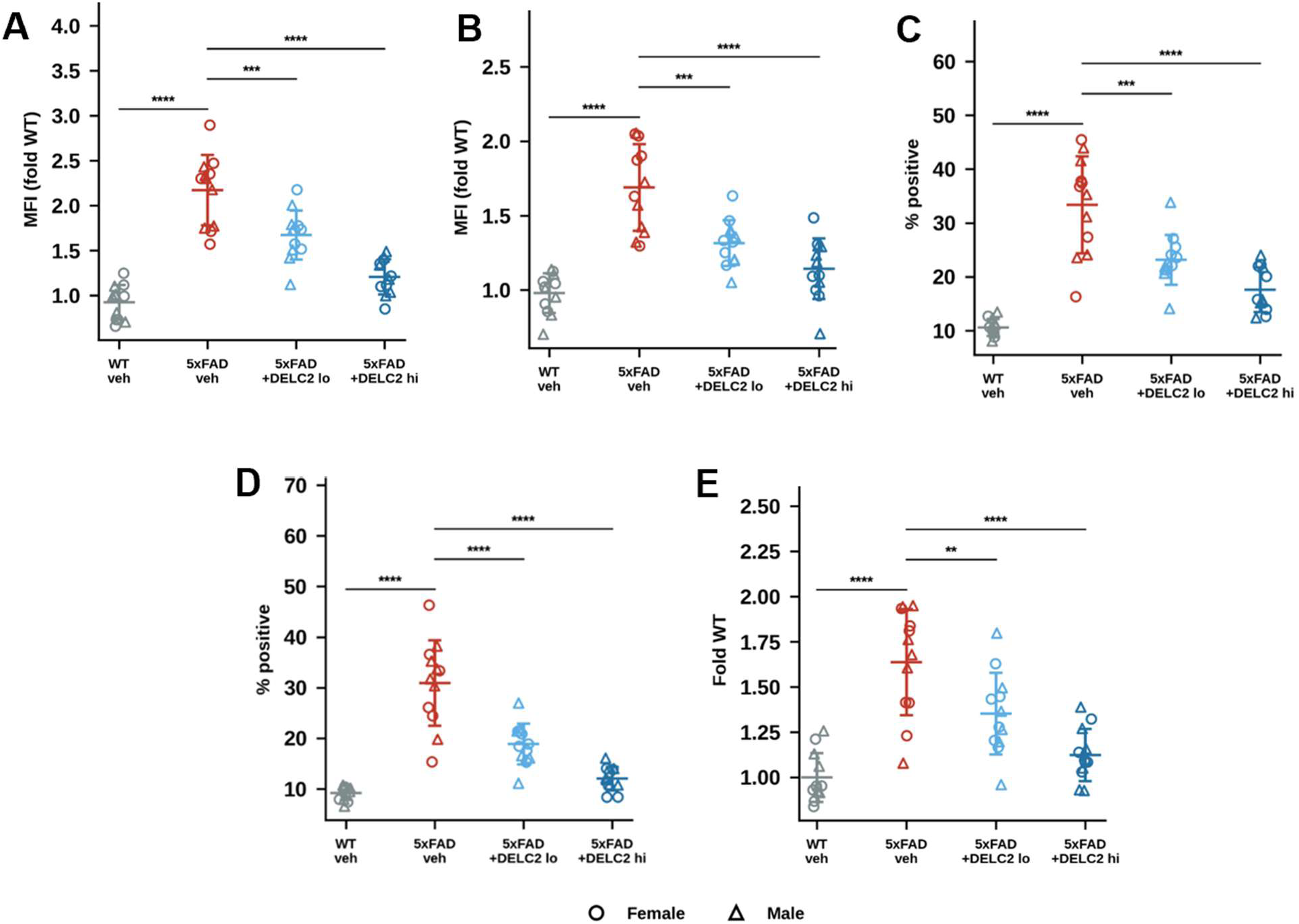
DELC2 normalizes cell-resolved glial inflammatory signaling in 5xFAD brain. Dissociated brain cells from WT vehicle, 5xFAD vehicle, and 5xFAD mice treated with low-(10 mg/kg) or high-dose (30 mg/kg) DELC2 were analyzed by flow cytometry. **(A)** Astrocytic pSTAT3 MFI, expressed relative to WT vehicle. **(B)** Astrocytic pNF-κB MFI, expressed relative to WT vehicle. **(C)** Percentage of Gal-3-positive microglia. **(D)** Percentage of pSTAT3-positive microglia. **(E)** Ager/RAGE signal in the analyzed brain-cell suspension, expressed relative to WT vehicle. Each symbol represents one mouse; circles and triangles denote female and male mice, respectively (n = 12 mice per group). Horizontal bars indicate mean ± SEM. Statistical significance is denoted as follows: ns, not significant; *p < 0.05; **p < 0.01; ***p < 0.001; and ****p < 0.0001.

### DELC2 normalizes inflammatory gene programs in FACS-purified glia

To define the cellular transcriptional programs associated with DELC2 treatment, astrocytes and microglia were FACS-purified from WT vehicle, 5xFAD vehicle, and DELC2-treated 5xFAD brains and analyzed by targeted RT–qPCR. In astrocytes, 5xFAD vehicle mice showed increased expression of *Chi3l1*, *Ager*, *Stat3*, *Socs3*, *Nfkbia*, *Lgals3*, and *C3* relative to WT controls (Figure 9). DELC2 treatment attenuated this astrocyte-associated inflammatory program, with the high-dose group showing the most consistent reduction in *Chi3l1*, *Ager*, *Stat3*, *Socs3*, *Nfkbia*, *C3*, and *Apoe*. These changes are consistent with reduced activation of the CHI3L1–RAGE-linked NF-κB/STAT3 axis in astrocytes.

Microglia from 5xFAD vehicle mice exhibited a distinct but overlapping inflammatory profile, characterized by increased expression of *Stat3*, *Lgals3*, *C3*, *Il1b*, *Tnf*, *Trem2*, and *Apoe* (Figure 9). DELC2 reduced multiple microglial inflammatory and disease-associated genes, including *Lgals3*, *Il1b*, *Tnf*, *Trem2*, and *Apoe*, with more substantial normalization in the high-dose group. Expression of *Chi3l1*, *Ager*, *Stat3*, *Socs3*, and *Nfkbia* was also reduced in microglia, supporting convergence of astrocyte and microglial responses on a shared CHI3L1-RAGE-associated signaling network. Together, these cell-type-resolved analyses show that DELC2 is associated with coordinated suppression of reactive astrocyte and inflammatory microglial gene programs in 5xFAD brain.

**Figure 9.**
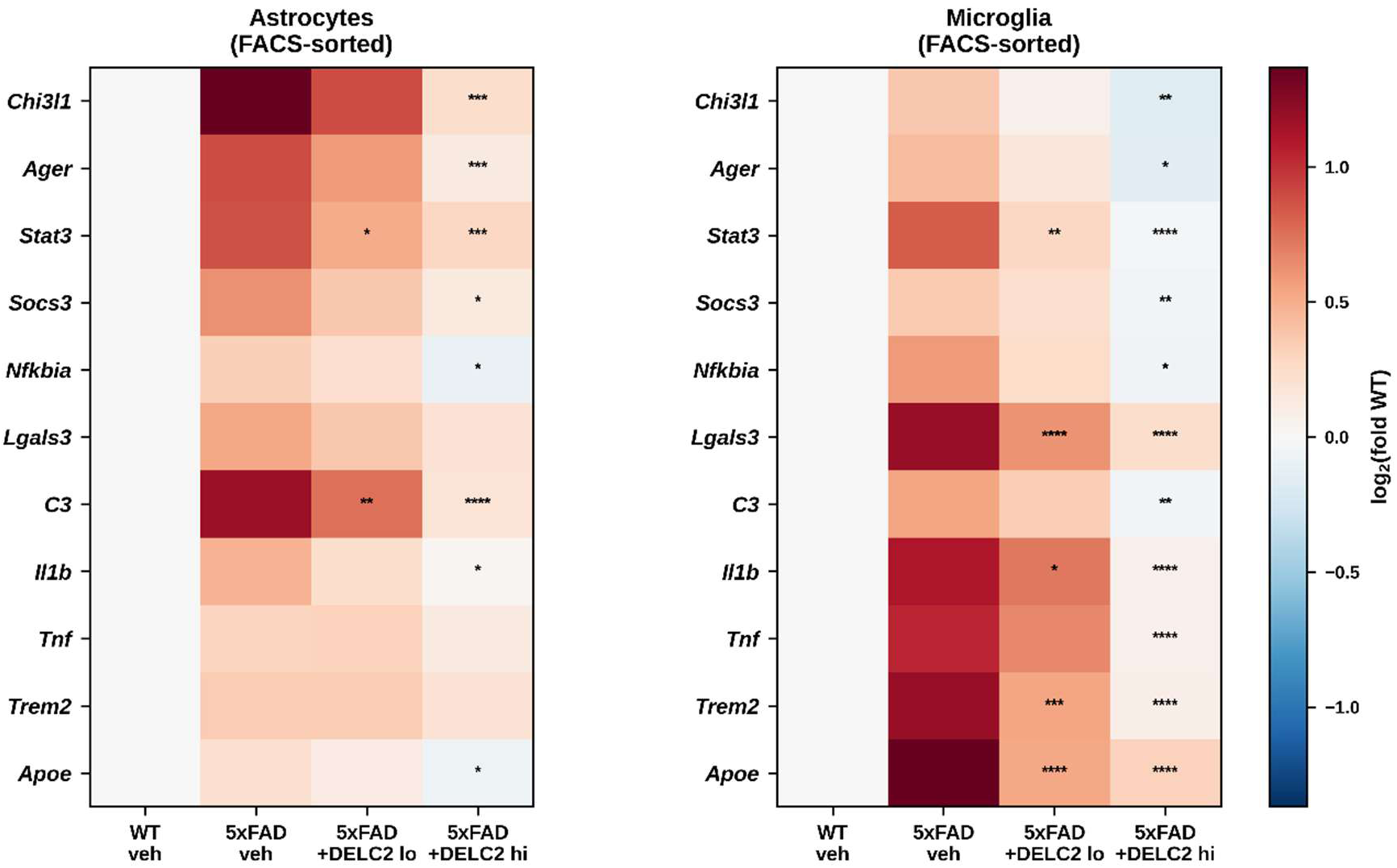
DELC2 normalizes inflammatory gene-expression programs in FACS-purified astrocytes and microglia from 5xFAD brain. RT-qPCR was performed in FACS-purified astrocytes and microglia isolated from WT vehicle, 5xFAD vehicle, and low-or high-dose DELC2-treated 5xFAD mice. Heatmaps show log_2_ fold change relative to the WT vehicle mean for each gene within each purified cell population. Asterisks indicate Tukey-adjusted comparisons with the 5xFAD vehicle group within the corresponding cell type. Statistical significance is denoted as follows: ns, not significant; *p < 0.05; **p < 0.01; ***p < 0.001; and ****p < 0.0001.

## Discussion

CHI3L1 is increasingly recognized as a prominent feature of astrocyte reactivity in Alzheimer’s disease, with elevated CSF levels associated with tau pathology, neurodegeneration, and cognitive decline (34). Genetic studies showing that astrocyte-specific deletion of *Chi3l1* reduces Aβ burden, glial activation, and cognitive impairment in 5xFAD mice further suggest that CHI3L1 contributes functionally to disease progression rather than serving solely as a biomarker (14). However, the molecular interactions through which CHI3L1 propagates glial inflammation and whether this extracellular pseudoenzyme can be pharmacologically targeted in the brain have remained poorly defined (35). In this study, we show that DELC2, a brain-penetrant small molecule identified by DNA-encoded library screening, engages the glycan-recognition groove of CHI3L1 and reduces its association with RAGE. DELC2 suppressed CHI3L1-driven inflammatory signaling, restored impaired Aβ handling in glial systems, attenuated the inflammatory activity of human AD CSF, and improved cognitive, amyloid-associated, and glial inflammatory outcomes following oral administration in 5xFAD mice. These findings extend previous genetic evidence by demonstrating that structurally defined engagement of CHI3L1 can pharmacologically modulate RAGE-linked neuroimmune dysfunction in AD.

The CHI3L1-DELC2 structure provides a molecular framework for targeting a chitinase-like protein that lacks measurable enzymatic activity and a conventional catalytic site (36). DELC2 occupies the canonical glycan-recognition groove within an aromatic-hydrophobic environment centered on Trp99 and Trp352. Rather than acting as a classical carbohydrate mimic, DELC2 achieves extended hydrophobic occupancy and shape-complementary packing across the recognition cleft. Because Trp99 is conformationally dynamic and contributes to ligand accommodation, stabilization of the surrounding aromatic network by DELC2 may restrict groove conformations required for physiological ligand or receptor interactions (27). Direct measurements of ligand-induced conformational dynamics will be required to test this possibility. Nevertheless, the structure demonstrates that the CHI3L1 recognition groove can accommodate a chemically distinct, drug-like ligand and provides a basis for structure-guided optimization of this compound class.

A central mechanistic finding was that DELC2 reduced CHI3L1-RAGE association in two independent biochemical formats. Consistent with this effect, DELC2 and RAGE blockade produced overlapping reductions in CHI3L1-induced NF-κB, IL-6, and STAT3 signaling. Both interventions also restrained the emergence of Gal-3-positive microglia and restored Aβ uptake in the astrocyte–microglia co-culture system. The limited additional effect of combining DELC2 with RAGE blockade on several endpoints supports functional convergence on a common pathway, although it does not establish complete mechanistic equivalence. Direct CHI3L1 neutralization produced effects in the same direction, providing an independent pathway control. Together, these results support a model in which extracellular CHI3L1 reinforces reactive glial states through RAGE-linked signaling, thereby coupling astrocyte-derived inflammatory activity to microglial dysfunction and impaired Aβ handling. Exploratory docking recurrently positioned a putative RAGE-contact region near the DELC2-binding groove, providing a possible explanation for the competition results. However, because the docking was unrestrained and did not converge uniformly, it should be considered hypothesis-generating. Structural determination of the CHI3L1-RAGE complex and mutational analysis of the proposed interface will be needed to distinguish direct steric competition from ligand-induced conformational modulation. Moreover, CHI3L1 can interact with additional receptors, including IL-13Rα2 and CD44, and the present results do not establish that all DELC2 effects are mediated exclusively through RAGE (37, 38).

The human CSF experiments strengthen the disease relevance of this pathway. CHI3L1 abundance correlated with the magnitude of astrocyte NF-κB activation induced by individual CSF samples. More importantly, paired immunodepletion and add-back experiments showed that removal of CHI3L1 attenuated inflammatory signaling and restored Aβ uptake, whereas reconstitution with recombinant CHI3L1 reinstated these phenotypes. DELC2 reproduced the direction of CHI3L1 depletion across astrocyte NF-κB activation, astrocytic Aβ uptake, and microglial IL-1β production. Although AD CSF contains numerous inflammatory and disease-associated mediators, the depletion-reconstitution design provides evidence that CHI3L1 contributes directly to its biological activity in this experimental system. These findings support CHI3L1 as an active and pharmacologically modifiable component of the AD-associated extracellular environment rather than solely a correlate of reactive gliosis.

Oral DELC2 administration produced sustained unbound brain exposure and improved performance in two complementary cognitive tasks in 5xFAD mice without altering exploratory activity. These behavioral effects were accompanied by reductions in soluble, insoluble, and oligomeric Aβ measures and partial normalization of the hippocampal Aβ42/Aβ40 ratio. DELC2 also reduced bulk-brain inflammatory measures and suppressed astrocytic and microglial NF-κB/STAT3-associated signaling. Targeted transcriptional analysis of FACS-purified glia extended these observations by showing coordinated attenuation of reactive astrocyte and inflammatory microglial gene programs. The convergence of behavioral, biochemical, cellular, and transcriptional responses is consistent with a mechanism in which CHI3L1 modulation improves glial homeostasis and Aβ handling rather than affecting a single inflammatory endpoint. The reduction in brain *Chi3l1* expression following treatment may reflect interruption of a feed-forward circuit in which CHI3L1-dependent signaling reinforces its production and maintains reactive glial states, although the regulatory basis of this response remains to be established.

While these findings link pharmacological CHI3L1 engagement to suppression of RAGE-linked glial inflammation, there are limitations to the present study. The biochemical competition and RAGE-neutralization experiments do not establish whether DELC2 directly occludes the RAGE-binding interface, alters CHI3L1 conformation, or affects its interactions with other receptors. Direct brain target occupancy was not measured, and the requirement for *Chi3l1* in the in vivo response to DELC2 remains to be established. Testing DELC2 in *Chi3l1*-deficient or astrocyte-specific *Chi3l1*-knockout AD models would provide a stringent assessment of target dependence, while pocket-directed mutagenesis could determine whether the crystallographically identified interactions are required for DELC2 activity.

It will also be important to determine whether the effects observed in 5xFAD mice extend to models incorporating tau pathology, synaptic degeneration, and the heterogeneity of sporadic AD. Although DELC2 reduced several Aβ measures, the current experiments do not distinguish altered production, uptake, degradation, aggregation, or distribution, and quantitative plaque histology was not performed. Longer treatment studies incorporating neuropathological, synaptic, pharmacodynamic, and chronic-safety endpoints will therefore be required. Validation in larger, clinically characterized CSF cohorts and human iPSC-derived glial systems should further establish the relevance of this pathway across patient populations and disease stages. Finally, DELC2 remains an early-stage scaffold with low-micromolar affinity, providing an opportunity for structure-guided optimization of potency, brain exposure, and target engagement. Together, the present findings establish the CHI3L1 glycan-recognition groove as a druggable regulatory site and support further development of brain-penetrant CHI3L1 modulators for AD and related neurodegenerative disorders.

## Conclusion

This study establishes CHI3L1 as a druggable regulator of RAGE-linked glial dysfunction in AD and identifies DELC2 as a brain-penetrant chemical scaffold for modulating this pathway. By combining structural target engagement, biochemical competition, mechanistic glial studies, human AD CSF experiments, and oral efficacy in 5xFAD mice, our findings connect occupancy of the CHI3L1 glycan-recognition groove with suppression of neuroinflammatory signaling, restoration of Aβ handling, and improvement of disease-associated outcomes. These results reposition CHI3L1 from a marker of reactive astrocytes to a pharmacologically actionable component of AD pathobiology and provide a foundation for developing more potent brain-penetrant CHI3L1 modulators for AD and related neurodegenerative disorders.

## List of abbreviations

AD: Alzheimer’s disease
ADME: Absorption, distribution, metabolism, and excretion
Aβ: Amyloid-β
BLI: Biolayer interferometry
CHI3L1: Chitinase-3-like protein 1
CSF: Cerebrospinal fluid
DEL: DNA-encoded library
FACS: Fluorescence-activated cell sorting
MST: Microscale thermophoresis
NF-κB: Nuclear factor kappa B
PK: Pharmacokinetics
RAGE: Receptor for advanced glycation end products
RT-qPCR: Reverse transcription quantitative polymerase chain reaction
SPR: Surface plasmon resonance
STAT3: Signal transducer and activator of transcription 3

## Data availability

All data needed to evaluate the conclusions in the paper are present in the paper and/or the Supplementary Materials. Atomic coordinates and structure factors for the CHI3L1-DELC2 complex have been deposited in the Protein Data Bank under accession code 10KA. All other data supporting the findings of this study are available within the article and its Supplementary Information files. The amino acid sequence of human CHI3L1 can be accessed through the UniProt Knowledgebase under UniProt ID P36222.

## Ethics and Consent to Participate declarations

not applicable

## Author contributions

Conceptualization, S.U. and M.T.G.; methodology, S.U., L.Z., B.K., L.C.B., M.T., and M.T.G.; investigation, S.U., L.Z., B.K., and L.C.B.; formal analysis, S.U., L.Z., B.K., L.C.B., and M.T.; structural biology and crystallographic analysis, S.U. and M.T.; resources, M.T. and M.T.G.; supervision, M.T.G.; project administration, M.T.G.; writing -original draft, S.U. and M.T.G.; writing -review and editing, S.U., L.Z., B.K., L.C.B., M.T., and M.T.G.

## Acknowledgement

This research used FMX beamline (17-ID-2) of the National Synchrotron Light Source II, a U.S. Department of Energy (DOE) Office of Science User Facility operated for the DOE Office of Science by Brookhaven National Laboratory under Contract No. DE-SC0012704. The Center for BioMolecular Structure (CBMS) is primarily supported by the National Institutes of Health, National Institute of General Medical Sciences (NIGMS) through a Center Core P30 Grant (P30GM133893), and by the DOE Office of Biological and Environmental Research (KP1605010).

