## Supporting Information for "Discovery of a brain-penetrant CHI3L1 inhibitor as an oral modulator of neuroinflammation in Alzheimer’s disease"

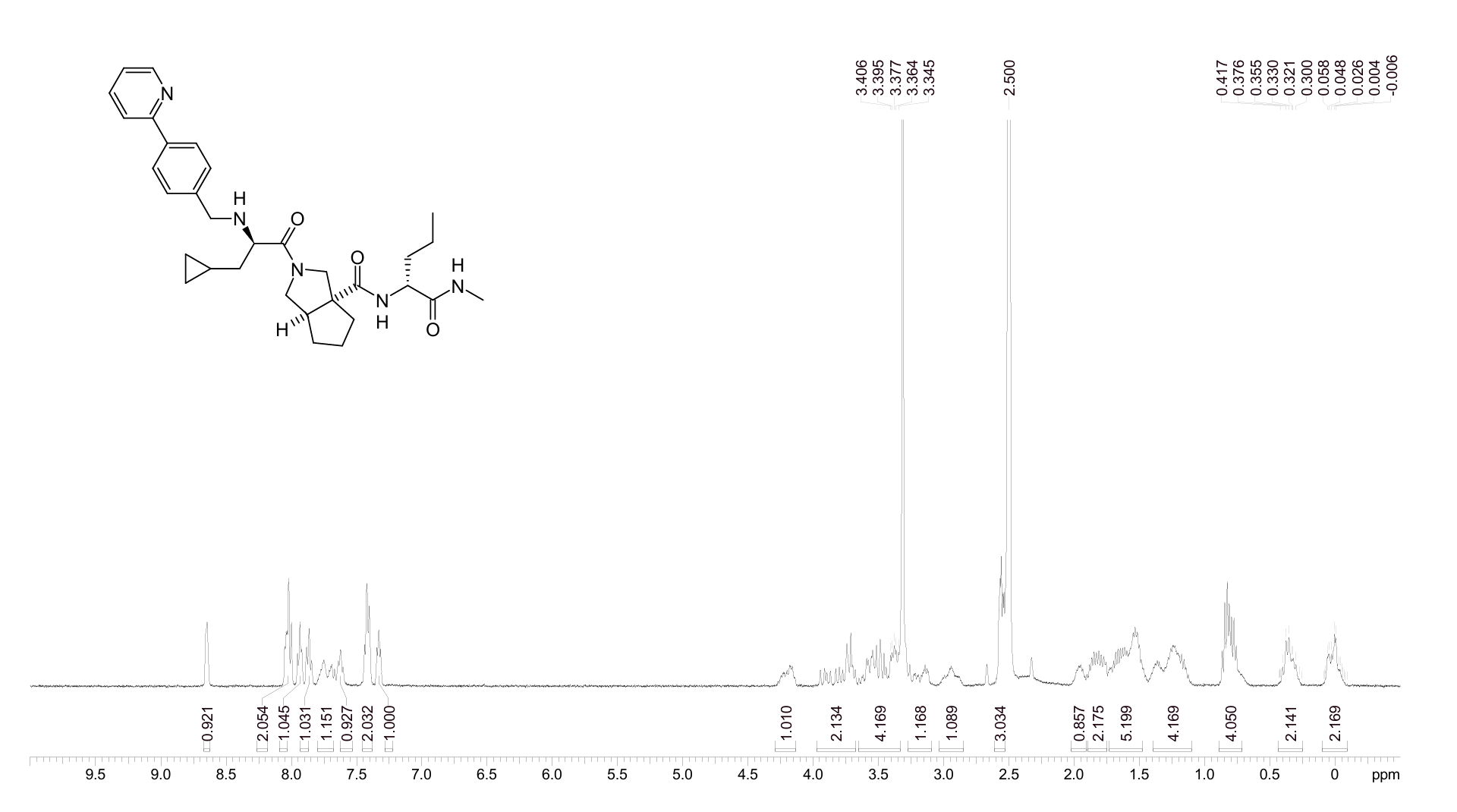

**Figure S1.** ^1^H NMR spectrum for DELC2.

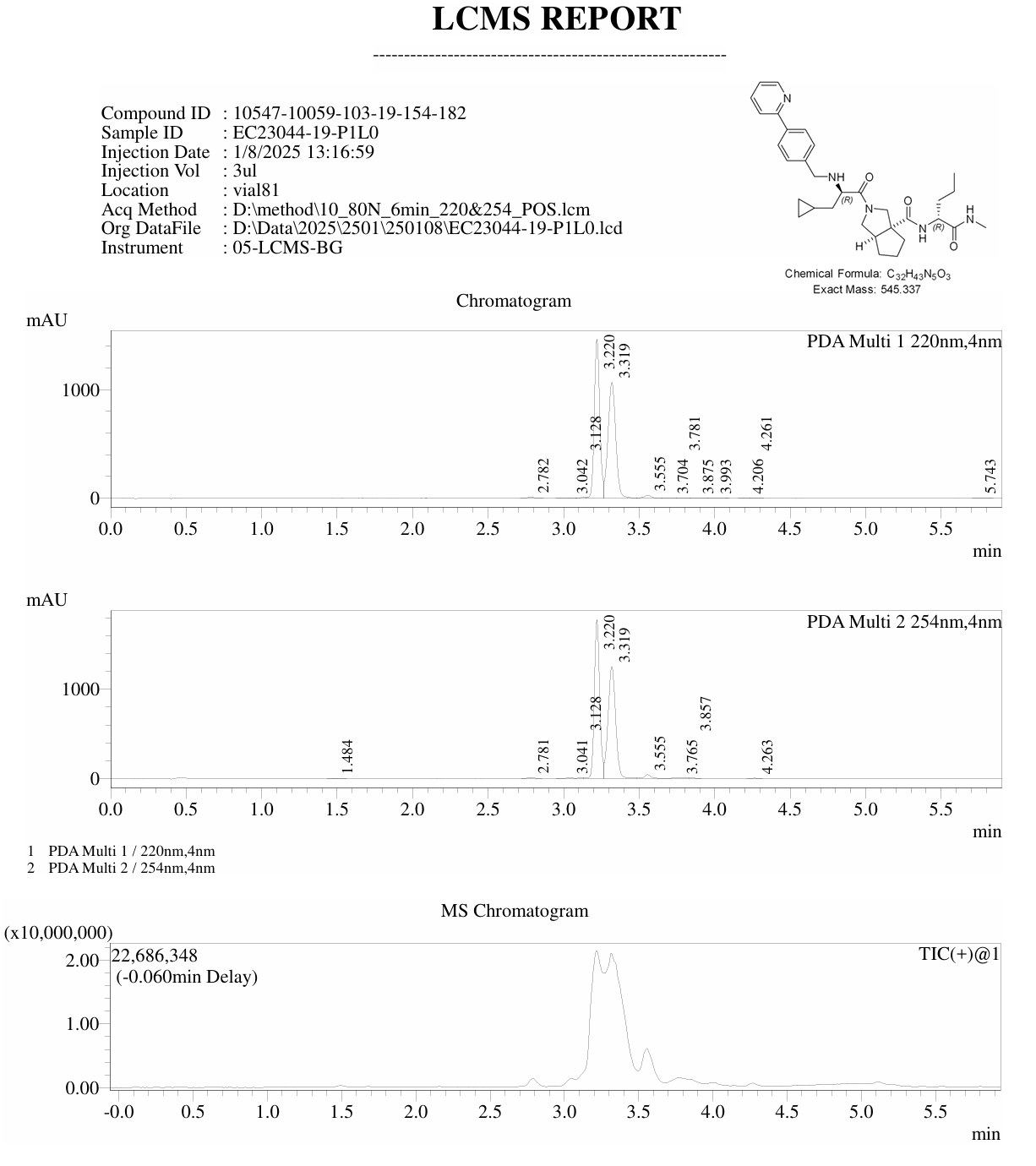

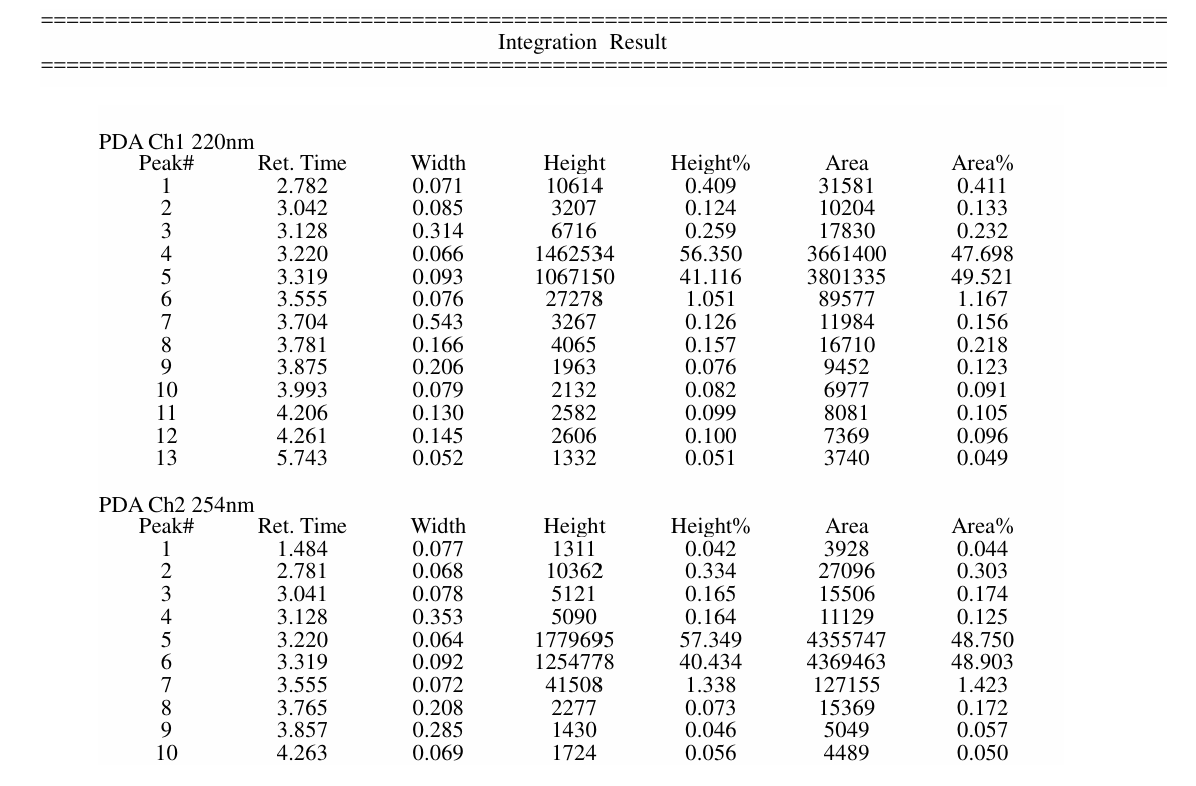

**Figure S2.** LCMS report for DELC2.

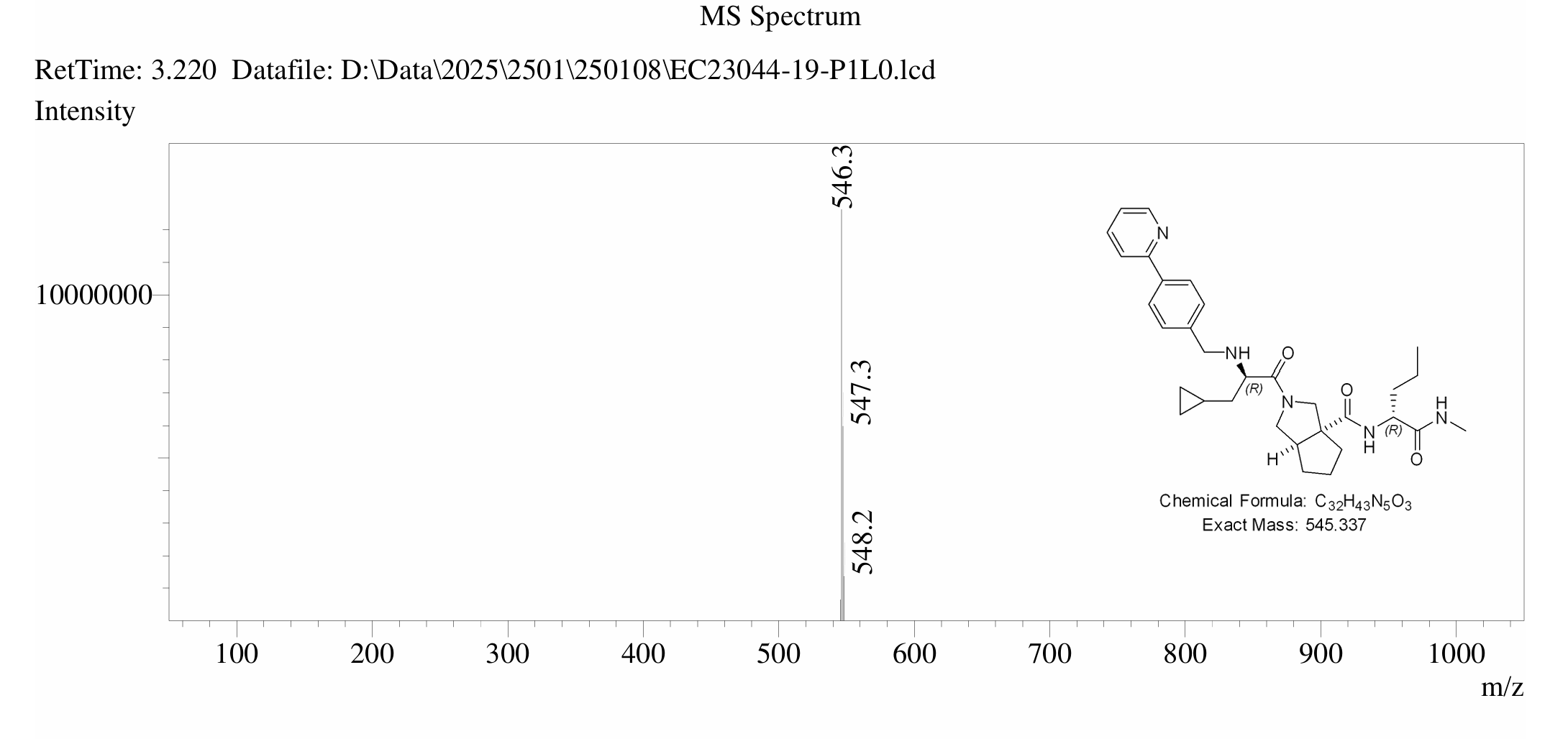

**Figure S3.** MS spectrum for DELC2.

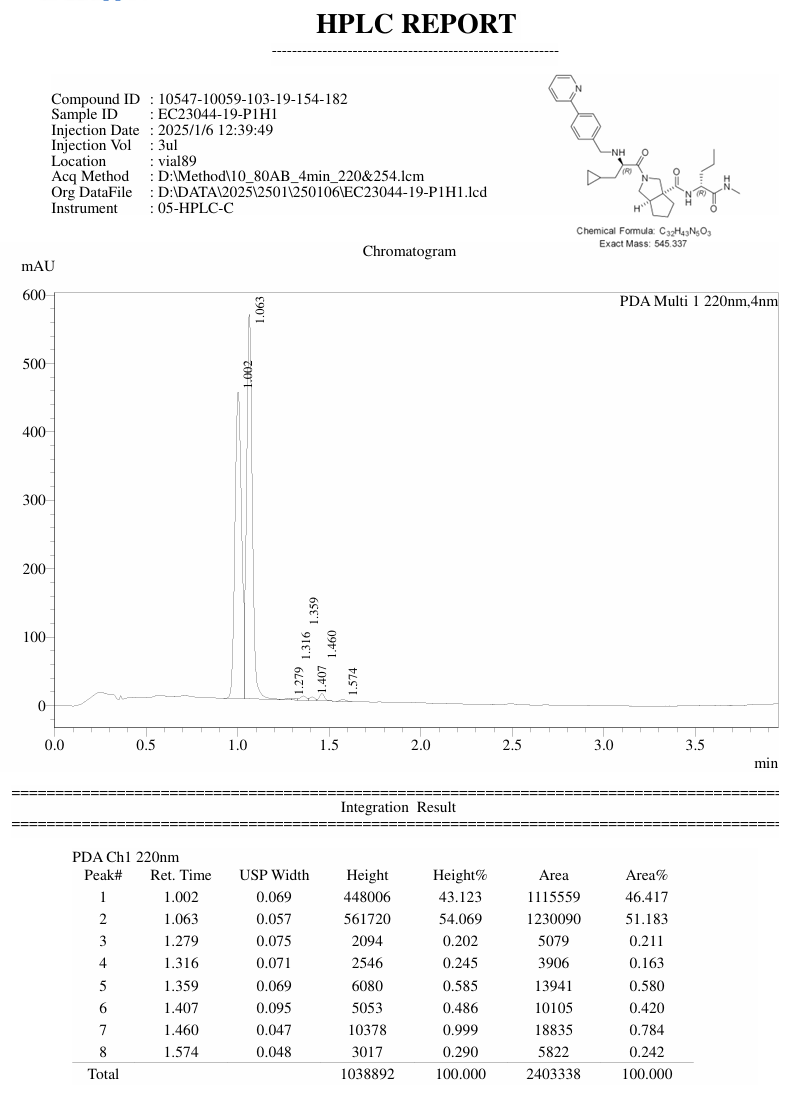

**Figure S4.** HPLC trace for DELC2.

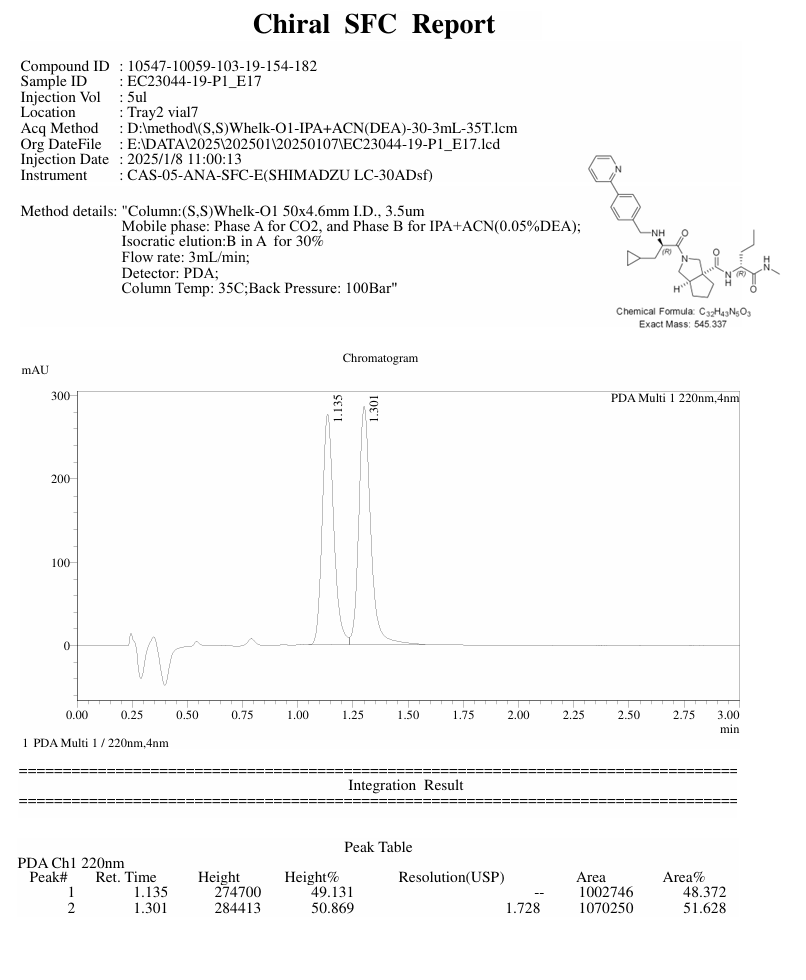

**Figure S5.** Chiral SFC report for DELC2.

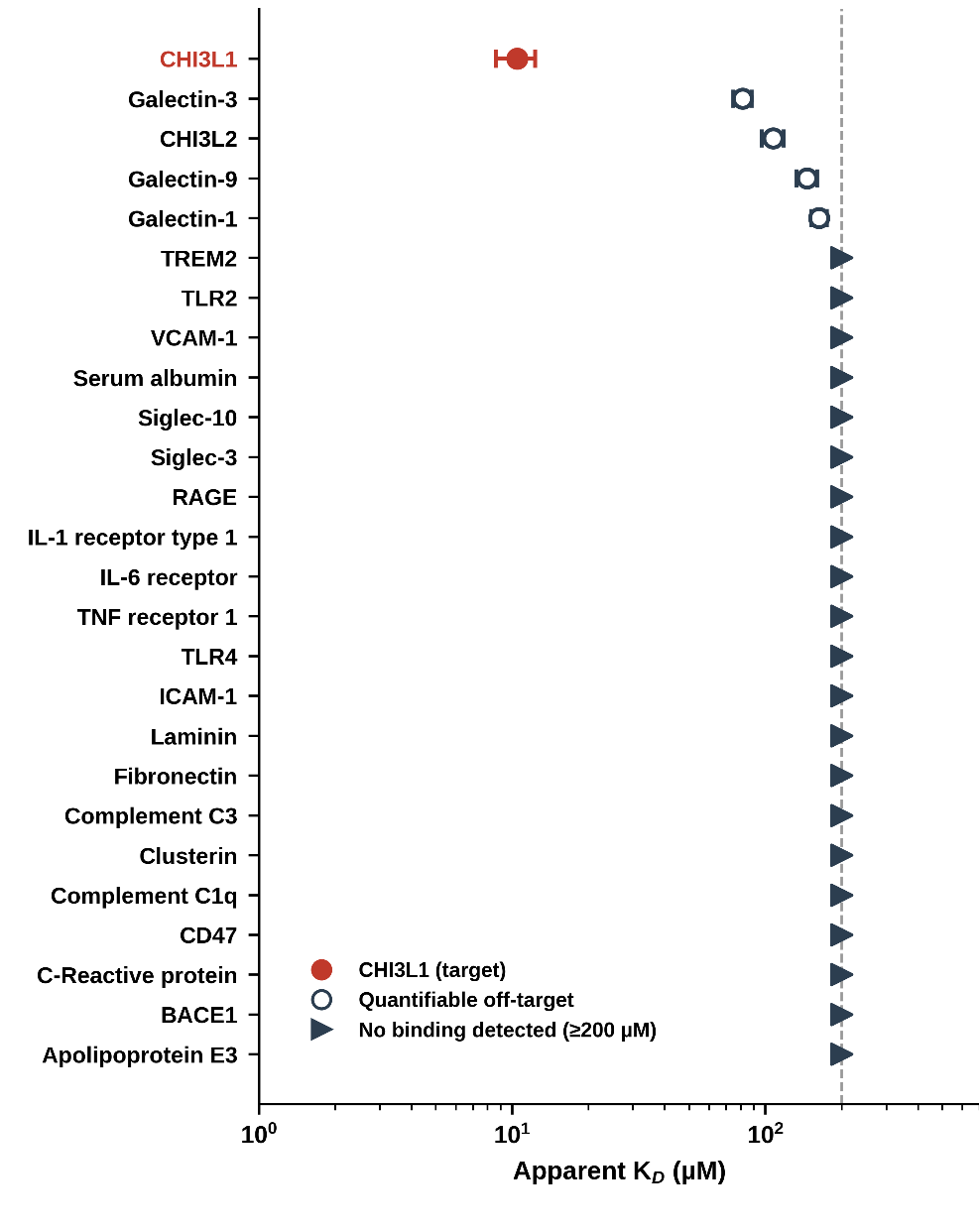

**Figure S6. MST counterscreen selectivity profile of DELC2 across a 25-protein panel.** Apparent dissociation constants (K_D_) are shown on a logarithmic scale. CHI3L1 is highlighted in red; open circles indicate proteins with quantifiable weak binding, and right-facing triangles indicate no quantifiable binding within the tested concentration range (0-200 µM).

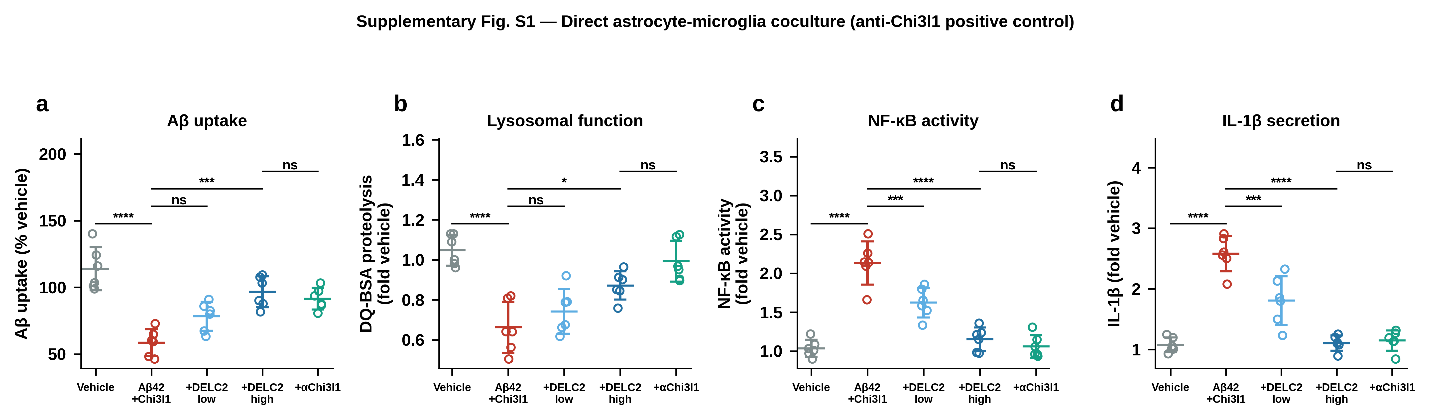

**Figure S7. Direct CHI3L1 neutralization phenocopies DELC2 in an astrocyte-microglia co-culture system.** Primary astrocyte-microglia co-cultures were exposed to Aβ_42_ and recombinant CHI3L1 in the presence of vehicle, low or high DELC2, or a CHI3L1-neutralizing antibody (αCHI3L1). **(A)** Microglial Aβ uptake. **(B)** Lysosomal proteolytic activity measured by DQ-BSA processing. **(C)** NF-κB activity. **(D)** IL-1β secretion. Individual points represent independent experimental replicates; horizontal bars indicate mean ± SEM (n = 5 independent biological replicates per condition). Statistical significance is denoted as follows: ns, not significant; *p < 0.05; ***p < 0.001; and ****p < 0.0001.

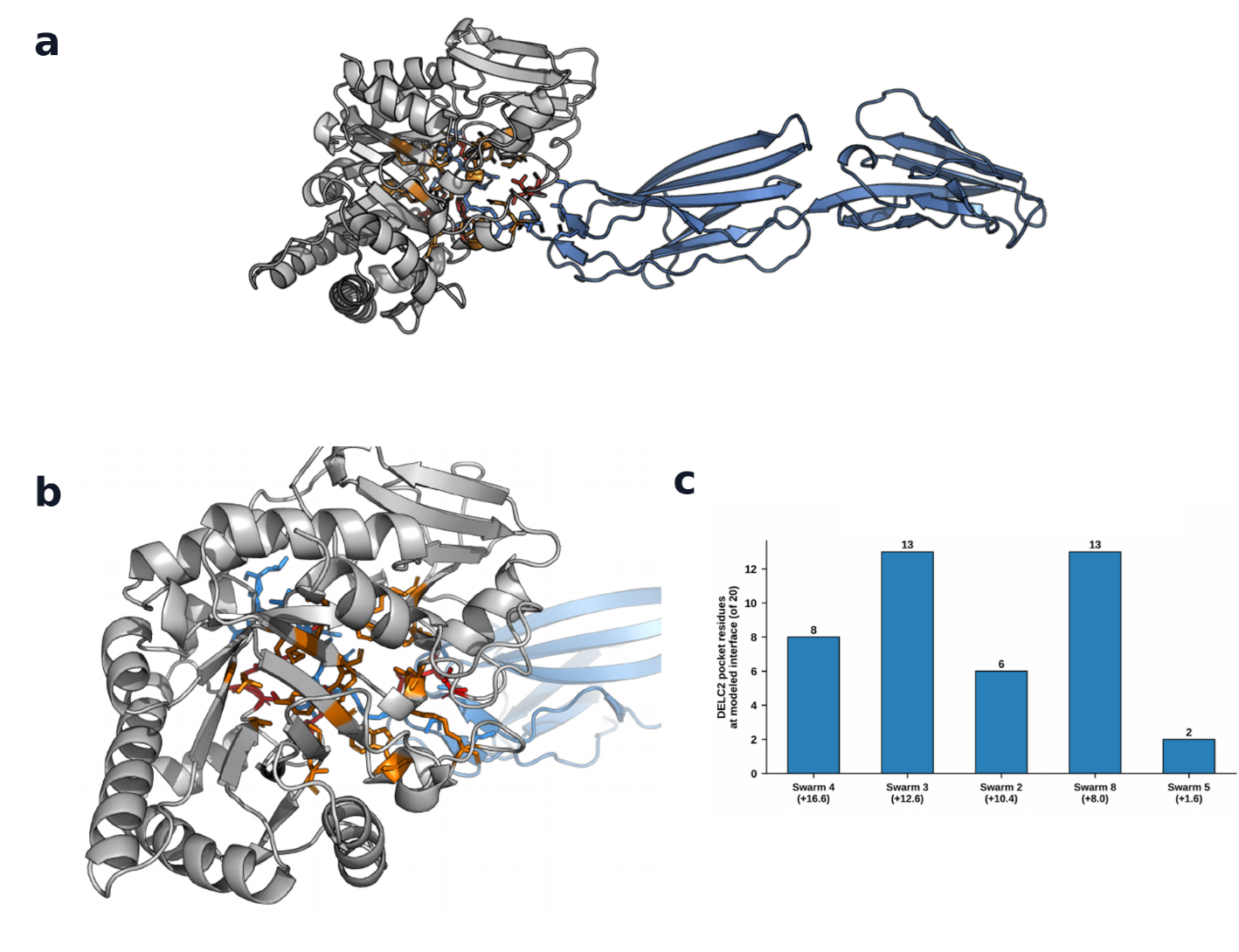

**Figure S8. Docking of RAGE near the CHI3L1 DELC2-binding groove. (A)** Representative full rigid-body docking pose of the RAGE V-C1 domain (blue) relative to CHI3L1 (gray). Residues comprising the crystallographically defined DELC2-binding groove are shown in orange; residues recurrently assigned to the modeled interface are highlighted in red. **(B)** Enlarged view of the modeled interface at the CHI3L1 glycan-recognition groove. **(C)** Number of DELC2-pocket residues located at the modeled RAGE-contact surface across the five favorable-scoring LightDock swarms; values in parentheses indicate the top-pose DFIRE score for each swarm. This blind rigid-body docking analysis is hypothesis-generating and does not define the native CHI3L1-RAGE interface.

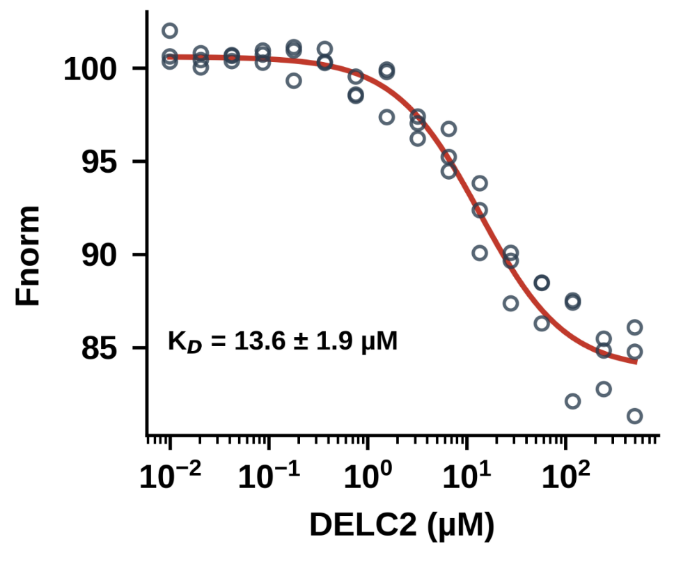

**Figure S9. DELC2 directly binds murine CHI3L1.** Microscale thermophoresis (MST) analysis of DELC2 binding to recombinant murine CHI3L1. Individual data points represent independent measurements (n = 3) across the DELC2 concentration series; the red line indicates the fitted binding curve.

**Table S1. Crystallographic Data Collection and Refinement Statistics for the CHI3L1-DELC2 Complex**

**Data Collection	DELC2(10KA)**

Space group	P 43 21 2

Cell dimensions

a, b, c (Å	104.02Å, 104.02Å, 161.68Å

α, β, γ (°)	90.00, 90.00, 90.00

Resolution (Å)	33.90 – 2.47

Wavelength (Å)	0.97934

Rmerge	0.29

I / σ(I) highest shell	0.57

Completeness (%)	100

Redundancy	17

**Refinement**

Resolution (Å)	33.90 - 2.47

No. reflections	32371

Rwork / Rfree	0.2194/0.2528

No. atoms

Protein	5706

Ligand / ion (DELC2+NAG)	136

Water	65

B-factors (Å²)

Protein	84.9

Ligand / ion	86.5

Water	58.0

R.m.s. deviations

Bond lengths (Å)	0.004

Bond angles (°)	0.814

**Table S2. In vitro absorption, distribution, metabolism, excretion, and early safety profile of DELC2.**

| **PK Parameter** | **DELC2** |
| --- | --- |
| LogD_7.4_ᵃ | 2.64 |
| Solubility, 1% DMSO/PBS [µM]ᵇ | 212 |
| Solubility in FaSSIF [µM] | 318 |
| PAMPA Pₐₚₚ [×10^-6^ cm/s] | 18.7 |
| Stability in simulated gastric fluid (pH 1.2, 2 h), % remaining | 98.8 |
| Stability in simulated intestinal fluid (pH 6.8, 2 h), % remaining | 95.3 |
| t½ [min], mouse liver microsomes | 75.3 |
| t½ [min], human liver microsomes | 92.5 |
| t½ [min], mouse plasma | >120ᶜ |
| t½ [min], human plasma | >120ᶜ |
| t½ [min], PBS | >120ᶜ |
| Human plasma protein binding [%] | 92.3 |
| IC_50_ against WI-38 [µM] | >100 |
| IC_50_ against HS-27 [µM] | >100 |
| IC_50_ against hERG [µM] | >100 |

ᵃ LogD at physiological pH. ᵇ Kinetic-solubility format. ᶜ The half-life was not reached within a 120-min assay window.
